# Tension TRAAKer: a chemigenetic fluorescent membrane tension reporter

**DOI:** 10.64898/2026.06.08.730921

**Authors:** Anna V. Elleman, Nels Gerstner, Benjamin E. Smith, Evan W. Miller, Richard H. Kramer, Stephen G. Brohawn

**Affiliations:** Department of Molecular and Cell Biology, University of California, Berkeley, CA, USA; Department of Neuroscience, University of California, Berkeley, CA, USA; California Institute for Quantitative Biosciences (QB3), University of California, Berkeley, CA, USA; Department of Chemistry, University of California, Berkeley, CA, USA

## Abstract

Mechanical force transduction is essential to survival, underlying biological processes as fundamental as morphogenesis, somatosensation, audition, and interoception; and driving pathologies as diverse as hypertension and cancer metastasis. Exogenous forces are translated to intracellular signals through transient changes in membrane tension which are currently not possible to directly monitor *in situ*. To remedy this, we have designed and validated Tension TRAAKer, a chemigenetic fluorescent membrane tension reporter for the visualization of tension induction, propagation, and dissipation in living cells. Tension TRAAKer is derived from inserting a tension-sensitive nonconductive variant of the mechanosensitive potassium ion channel TRAAK into a self-labelling HaloTag. Increasing membrane tensions effect conformational changes in the TRAAK channel that are optically monitored by a HaloTag-conjugated fluorogenic (environment-sensitive) dye. EGFP incorporation C-terminal to the HaloTag enables unambiguous tension reporting in mobile membranes via dual-color ratiometric imaging that controls for variations in sensor density. Tension TRAAKer reports membrane tension changes rapidly, reversibly, and with spatiotemporal precision—its fluorescence scaling to both stimulus magnitude and area, with consistent effect sizes observed between diverse cell types. It better distinguishes among elevated membrane tensions than do available indirect chemical reporters, with the additional advantage of being readily genetically targetable. We thus expect Tension TRAAKer to be a powerful tool for the study of membrane tension across biological systems and disease states.

## Introduction

The induction, maintenance, propagation, and dissipation of lipid membrane tension is fundamental to biological (dys)function, underlying diverse sensory processes like somatosensation, interoception, and audition^1^ and diseases including hypertension,^2^ cancer,^3^ and stroke.^4^ Membrane tension is central to cell differentiation,^5,6^ adhesion,^7^ proliferation,^8^ migration,^9,10^ and apoptosis,^11^ as well as the regulation of endo and exocytosis. ^12^ Reported basal membrane tensions are heterogeneous, ranging from 0.01 to 1 mN/m. ^13, 14, 15^ Cells integrate a variety of external stimuli that differentially alter membrane tension, including hydrostatic pressure, shear force, compressive stress,^16^ and chemical cues^17^ via a cohort of mechanosensitive channels that respond to tensions ranging from <1 mN/m (e.g., Piezo1) to >10 mN/m (e.g., MscL).^18^ Tension also propagates from the plasma membrane through the actin and microtubule cytoskeleton^19^ to intracellular organelles including the endoplasmic reticulum^20^ and nucleus^21^ via tether complexes.

Despite the importance of membrane tension to cell function, there are no existing tools that directly report membrane tension without perturbing physiology. While Flipper-TR® (Fluorescent LIPid Tension Reporter)^22^ is a widely-used membrane-inserting small molecule tension probe, it actually reports changes in lipid fluidity and composition, as its fluorescence lifetime varies according to lateral packing.^23^ Flipper-TR® poorly differentiates among basal and elevated membrane tensions, with tensions >0.5 mN/m, or osmolarities <300 mOsm, being functionally indistinguishable. Furthermore, as a chemical probe, Flipper-TR® nonspecifically labels all cells in a population. To address these gaps in current methodology, we aimed to design and optimize a generalizable fluorescent protein mechanosensor.

The effective monitoring of membrane tension requires a minimally perturbative tool with a rapid, reversible reaction to tension changes and a spatially precise, high fidelity, high signal to noise fluorescent output. Mechanosensitive ion channels meet a number of these requirements—they open rapidly (µs–ms) and reversibly in response to localized changes in membrane tension in a graded, reproducible manner. The incorporation of a conformation- or environment-sensitive fluorophore to an appropriately tension-sensitive location in these proteins might therefore be expected to yield a fluorescent mechanosensor. Other groups have successfully pursued a similar strategy to design fluorescent voltage sensors by taking advantage of the small (<5 Å) depolarization-induced transmembrane helix movements observed in certain voltage-dependent phosphatases (ASAP1,^24^ HArcLight).^25,26^

TWIK-related arachidonic acid-stimulated K^+^ channel (TRAAK) is a dimeric two-pore domain potassium channel (K_2P_) that is uniquely suited to the development of a fluorescent tension sensor. TRAAK (*Homo sapiens*) is small, with ∼285 amino acids (aa) within the transmembrane channel core; is activated by a broad range of tensions (10 to 90% activation range: 0.6–8.2 mN/m);^18^ and exhibits a structurally well-characterized, tension-induced conformational change. ^27, 28^ Each TRAAK monomer comprises four transmembrane helices (TMs), two pore helices, and a long (>100 aa) cytoplasmic, structurally unresolved, C-terminal tail.^29^ The two pairs of pore helices within a TRAAK dimer compose a pseudo four-fold symmetric G(Y/F)G potassium selectivity filter surrounded by TMs. Membrane tension favors lateral expansion of the TMs, most significantly in the TM4 C-termini, which are bent upward and outward by 15–25° (amounting to a 6–9 Å displacement) relative to the closed or leak open states.^28^ Channel opening to the mechano-activated state is largely voltage-insensitive, rapid (*τ*_open_ ∼0.23 ms),^30^ and fully reversible upon membrane relaxation.

Herein, we show that we can exploit this conformational change with the conjugation of a HaloTag and fluorogenic dye to create a fluorescent tension reporter. We further identify point mutations within the channel selectivity filter that ablate potassium conduction while maintaining tension sensitivity. In addition, by including a C-terminal EGFP in our sensor, we enable ratiometric tension monitoring even in mobile membranes. We call this reporter Tension TRAAKer.

### TRAAK HaloTag Design

To monitor TRAAK’s tension-induced TM4 conformational changes, we designed a suite of fluorescent mechanosensors. All rely on the positioning of a red fluorogenic rhodamine-based chloroalkane dye near the C-terminus of the TM4 helix via a self-labeling HaloTag (HT) protein (**Figure 1a**–**b**). The dye equilibrates between a non-fluorescent, cell permeable, lactone and a fluorescent zwitterion according to its local environment (**Figure 1b**–**d**). HaloTag-dye ligation, which forms a long-lasting ester bond, favors the fluorescent isomer, yielding fluorescence “turn-on”. Subsequent conformational shifts within, and movement of, the HaloTag—e.g., from TRAAK opening/closing, which conveys the HaloTag to/from the channel mouth and associated lipid membrane—are expected to alter the local environment of the dye and thus fluorescence output (**Figure 1a**–**c**).^25^ All sensors also include the full TRAAK C-terminus post-HaloTag, to ensure robust plasma membrane localization via TRAAK’s Ankyrin G binding sequence;^31,32^ followed by an EGFP protein, which enables dual color imaging to control for both channel expression and membrane movement. For initial screening experiments in HEK293T cells, each sensor terminated in a C-terminal CAAX farnesylation motif (denoted x in all constructs) to reduce fluorescence in internal lipid membranes (**Figure 1e-h**, see representative cells in **Figures 1**–**2** vs **3**–**6** for CAAX tag effects).^33,34^

**Figure 1:**
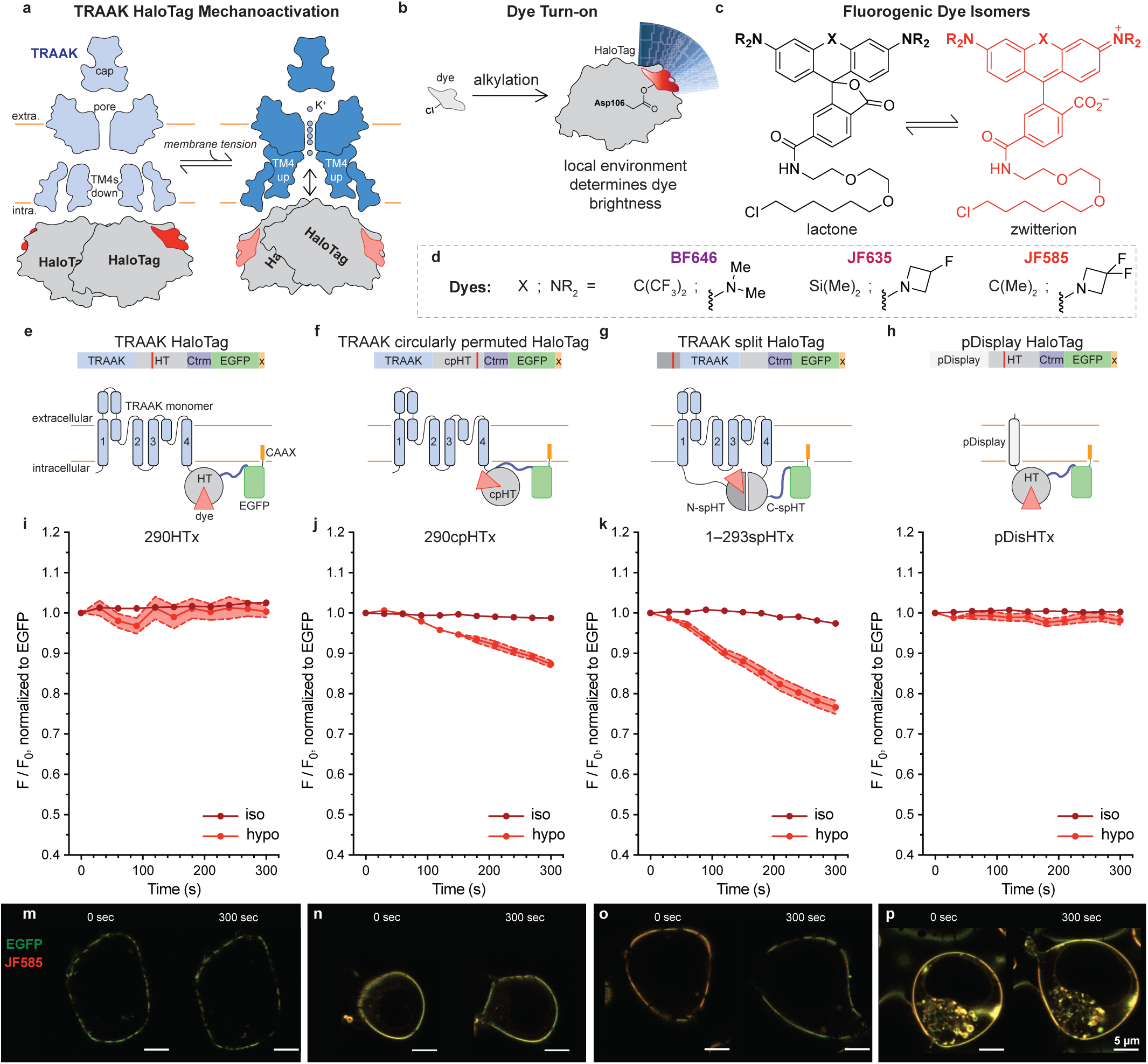
Design and validation of TRAAK/HaloTag-based membrane tension reporters. (**a**) Cartoon depiction of TRAAK HT dimer mechano-activated opening and concomitant TM4 upward/outward shift which reorients the HaloTag domain and attached dye. Extra./intra. = extracellular/intracellular. (**b**) Cartoon depiction of fluorogenic dye fluorescence change upon HaloTag Asp106 alkylation (turn on) and subsequent environment and/or conformational change about the dye binding site. Darker red = brighter dye; lighter red = dimmer dye. (**c**) Fluorogenic dye scaffold; X and NR_2_ defined in **d**. (**d**) Fluorogenic dyes. (**e**–**h**) Cartoon depictions of TRAAK HaloTag, TRAAK cpHaloTag, TRAAK spHaloTag, and pDisplay HaloTag protein sequences and structures, respectively. HT = HaloTag, cp = circularly permuted, sp = split, Ctrm = TRAAK C-terminus, x = CAAX. TRAAK core, blue; (cp/sp)HaloTag, grey; fluorogenic dye, red; Ctrm, purple; EGFP, green; CAAX tag, yellow. Transmembrane helices labeled 1–4. (**i**–**l**) Change in brightness of JF_585_ relative to EGFP over time in membranes of HEK293T cells transiently transfected with 290HTx, 290cpHTx, 1–293spHTx, or pDisHTx, respectively. F/F_0_ = dye:EGFP fluorescence at given time point / dye:EGFP fluorescence at t = 0 seconds. Iso/hypo: isotonic/hypotonic solution added. Data shown as mean ± s.e.m. n ≥ 21. (**m**–**p**) Images of representative cells from hypotonic condition presented at listed time points for 290HTx, 290cpHTx, 1–293spHTx, or pDisHTx, respectively. JF_585_ depicted in red, EGFP in green. Brightness scaling maintained within same channel and cell; scaling varies from cell to cell. Distance scale bar = 5 µm.

**Figure 2:**
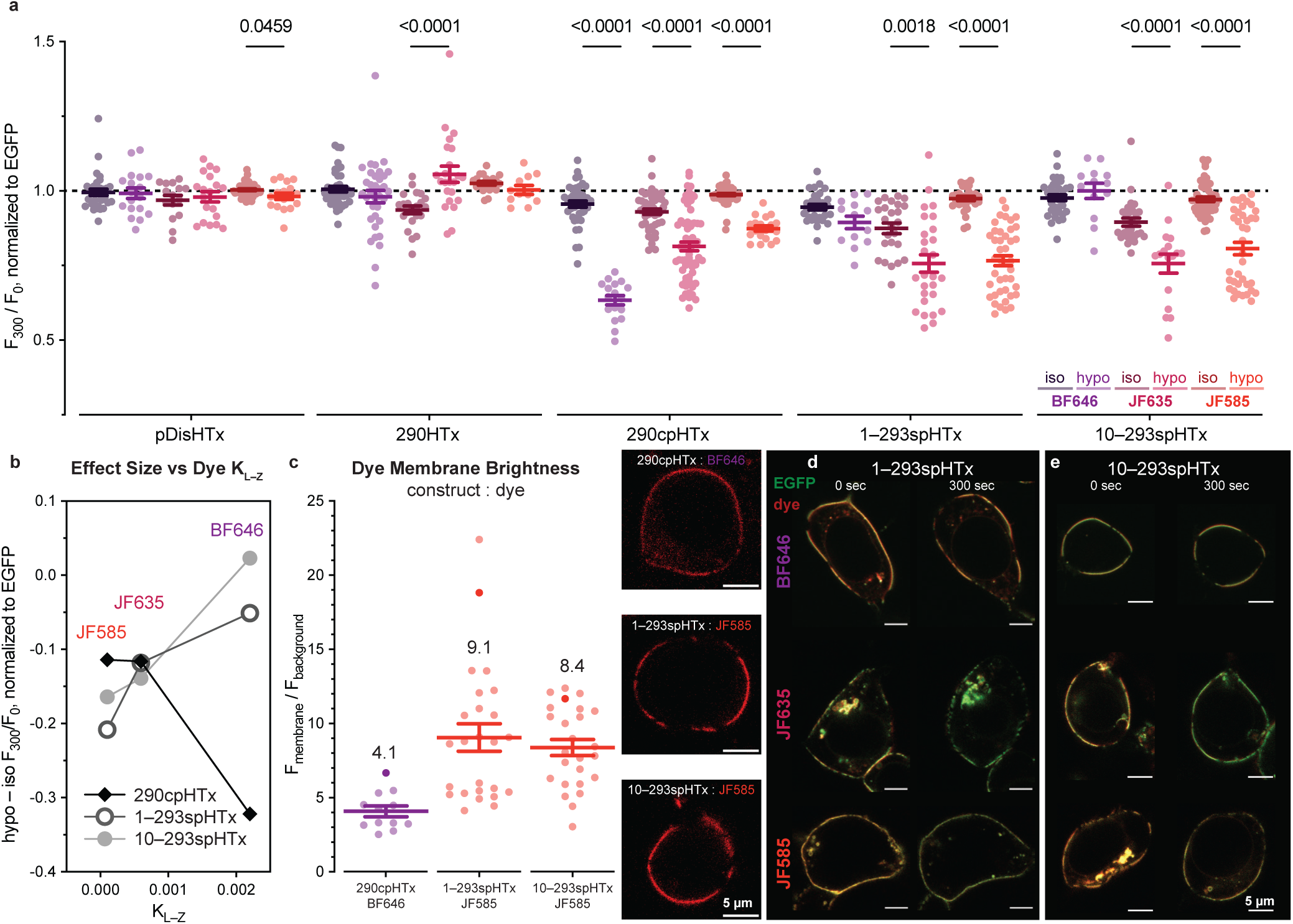
TRAAK (cp/sp)HaloTag hypotonic shock response varies according to fluorogenic dye identity. (**a**) Change in brightness of fluorogenic dye relative to EGFP after 5 minutes in membranes of HEK293T cells transiently transfected with described constructs. Iso/hypo: isotonic/hypotonic solution added. F_300_/F_0_ = dye:EGFP fluorescence at t = 300 seconds / t = 0 seconds. Data shown as mean ± s.e.m. with individual n’s as circles. n ≥ 13. p values calculated from multiple unpaired Mann-Whitney tests. p values < 0.05 provided above related data sets. (**b**) Hypo – Iso F_300_/F_0_ (normalized to EGFP) plotted against published dye K_L–Z_ values. (**c**) Relative dye membrane to background brightness of most responsive construct:dye pairs. Mean values provided above data. Representative membranes to right (dye in red, EGFP overlay omitted for clarity). Data points corresponding to representative membranes shaded darker. (**d**–**e**) Images of representative cells from hypo condition presented at listed time points after treatment with either BF_646_, JF_635_, or JF_585_ for 1–293spHTx, or 10–293spHTx, respectively. Fluorogenic dye depicted in red, EGFP in green. Brightness scaling maintained within same channel and cell; scaling varies from cell to cell. Distance scale bar = 5 µm.

Sensor efficacy was assessed via confocal microscopy. A z-stack of fluorogenic chloroalkane dye-bathed, sensor-expressing HEK293T cells was collected every 30 seconds immediately following a 200 mOsm hypotonic shock to increase membrane tension (**Supplementary Figure 1a**). Dye fluorescence intensity was quantified relative to expressed EGFP, enabling precise masking of the plasma membrane as each cell distended, such that changes in sensor fluorescence could be normalized to density regardless of membrane movement (**Supplementary Figure 1b**).

### TRAAK HaloTag Screen

The first TRAAK HaloTag tension sensors (TRAAKHTxs) we examined were analogous to the published voltage sensor HArcLight,^25^ bearing a C-terminal HaloTag (HT) protein conjugated to the base of TRAAK’s TM4 helix (**Figure 1e**, **Supplementary Figure 2a**). Two sensors in this category were studied: 285HTx and 290HTx, each named for the final residue in the TM4 helix prior to the HaloTag. Neither 285HTx (**Supplementary Figure 3b**) nor 290HTx (**Figures 1i**, **1m**) exhibited a consistent change in Janelia Fluor® 585 chloroalkane dye (JF_585_) brightness in response to hypotonic shock over a period of 5 minutes. In light of these results, we considered whether the dye might be misoriented relative to TRAAK in the wildtype HaloTag isomer, wherein the dye binding site faces the cytoplasm. Indeed, AlphaFold predictions of TRAAKHTx structures (cartooned in **Supplementary Figure 2a**) suggest that the fluorogenic dye is oriented away from the mouth of the channel and may thus experience little change in environment upon TM4 movement.

To better orient the fluorogenic dye, we next constructed a TRAAK channel bearing a circularly permuted HaloTag (cpHT) comparable to that described in the calcium sensor HaloCaMP^25^ (**Figure 1f**). Relative to HT, cpHT is split 37 residues beyond the dye alkylation site and circularized with a pentameric GG(S/T) linker connecting the original N- and C-termini, which we reasoned would reposition the dye toward the channel pore (**Supplementary Figure 2b**). Unlike 290HTx, 290cpHTx responds well to hypotonic shock, with an 11% reduction in JF_585_ intensity observed over 5 minutes (**Figure 1j**, **n**) compared to the isotonic control.

Finally, to further constrain the fluorogenic dye, we constructed a third series of channels using a “split” HaloTag. In these constructs, TRAAK is inserted into the HaloTag at the previously described circular permutation site (amino acid 142), dividing the HaloTag primary sequence such that the N-terminal portion (N-spHT) carries the dye alkylation residue (Asp106) and the C-terminal portion (C-spHT) the bulk of the dye binding site (**Figure 1g**).^35^ We reasoned this would orient the dye toward the membrane, similar to 290cpHTx, but additionally restrict dye rotational freedom (**Supplementary Figure 2c**). To vary the distance between the dye and the channel body, TRAAK’s flexible N- and C-termini were treated as linkers to be serially truncated toward the transmembrane core. Split HaloTag TRAAKs 1–293spHTx, 10–293spHTx, and 10–312spHTx—named for the spHT-encompassed TRAAK residues—exhibit larger responses to hypotonic shock than 290cpHTx with a 21%, 16%, or 22% reduction in JF_585_:EGFP signal observed after 5 minutes, respectively (**Figure 1k**, **Supplementary Figures 3e**–**g**). We attribute this improved responsiveness to conformational and/or environmental changes about the dye binding site as TM4 moves.

Control experiments were performed with pDisHTx, which comprises a tension-insensitive single transmembrane helix, pDisplay,^36^ in place of the TRAAK channel body in construct 285HTx (**Figure 1h**, **Supplementary Figure 2d**). JF_585_:EGFP fluorescence remained unchanged in HEK293T cells expressing this construct under all of our experimental conditions (**Figure 1l**, **1p**, **Supplementary Figure 3a**).

### Fluorogenic Dye Screen

To both corroborate our hypothesized TRAAK(cp/sp)HTx mechanism of action and to ensure maximal hypotonic shock response, we compared three red fluorogenic dyes with differing spectrochemical properties: carborhodamine JF_585_, Si-rhodamine JF_635_,^37^ and bis(trifluoromethyl)carborhodamine BF_646_^38^ (**Figure 1c**–**d**). All three dyes brighten in response to HaloTag alkylation due to stabilization of the fluorescent zwitterion compared to the uncharged lactone: JF_585_ by 98-fold, JF_635_ by 66-fold,^37^ and BF_646_ by 23-fold^38^ (**Supplementary Table 1**). In solution, JF_585_ most favors the nonfluorescent lactone, with a lactone-zwitterion equilibrium constant (K_L–Z_) of 0.0001,^39^ compared to 0.0006 for JF_635_^40^ and 0.0022 for BF_646_.^38^ JF_585_ is also the brightest of the three dyes, followed by JF_635_ and then BF_646_ (χχπ = 121680 vs 93520 vs 19140 M^−1^cm^−1^). We expected functional TRAAK(cp/sp)HTx constructs to exhibit different hypotonic shock-elicited changes in brightness according to bound dye, but the pDisHTx control and nonfunctional sensors to be agnostic to dye identity.

In accordance with our prediction, pDisHTx control experiments showed only minor hypotonic shock-induced change in red fluorescence irrespective of dye identity (**Figure 2a**, far left, maximum effect size = 2.1%, JF_585_). Tension insensitive TRAAKHTx constructs 285HTx (**Supplementary Figures 3b**, **4b**, **5b**) and 290HTx (**Figure 2a**, middle left) showed similarly negligible hypotonic shock-induced changes in brightness with all dyes. Construct 290cpHTx, however, exhibited a much larger response to hypotonic shock when bound to BF_646_ than JF_635_ or JF_585_: iso_300_–hypo_300_ = 32.2% vs 11.6% vs 11.4%, respectively (**Figure 2a**, middle). 290cpHTx’s responsiveness to tension was thus correlated with the solution K_L–Z_ of the conjugated fluorogenic dye, with dimmer, poorer “turn-on” dyes yielding larger responses (**Figure 2b**, **Supplementary Table 1**).

TRAAKspHTx constructs 1–293spHTx and 10–293spHTx followed the opposite trend and were most responsive when conjugated to lower K_L–Z_, brighter, better turn-on dyes (**Figure 2a**, right; **2b**, **2d**–**e**): e.g., 1–293spHTx, iso_300_–hypo_300_ = 5.1% vs 11.8% vs 20.8%, for BF_646_, JF_635_, and JF_585_ respectively. We suspect this may arise from partial re-solvation of the dye binding site under tension. Both constructs exhibited some degree of photobleaching when conjugated to JF_635_ (see untreated vs iso conditions, **Supplementary Figures 4f**–**g**), rendering precise quantification of hypotonic shock effect size difficult. In contrast, the longest TRAAKspHTx construct, 10–312spHTx, did not respond to hypotonic shock at all when conjugated to JF_635_ (**Supplementary Figures 3e**, **4e**), reaffirming that tight connection between the base of the TM4 helix and the C-terminal spHT domain improves the tension response. Complete time course data for all constructs and dyes are provided in **Supplemental Figures 3–6**.

To evaluate reporter practicality, we examined the membrane dye brightness of the most responsive TRAAK(cp/sp)HTx:dye pairs—290cpHTx:BF_646_, 1–293spHTx:JF_585_, 10–293spHTx:JF_585_ (**Figure 2a**) relative to background. 290cpHTx:BF_646_ offered substantially worse signal to noise than either spHT:JF_585_ pair, with dye signal within the plasma membrane averaging only ∼4x above background (**Figure 2c**) even before hypotonic shock-induced dimming. In comparison, spHTx:JF_585_ pairs produced plasma membrane dye signals 8–9x brighter than background, ranging from 4–22x in the case of 1–293spHTx:JF_585_. Note that no dye wash-out was performed here, or in any of the other osmolarity experiments herein, as one of the key advantages of a turn-on dye is its low background absent wash. We attribute 290cpHTx:BF_646_’s dimness relative to the spHTx:JF_585_ pairings to differences in construct expression efficacies, dye binding site environments, and dye spectrochemical properties (lower brightness and fold turn-on of BF_646_ vs JF_585_, **Supplementary Table 1**). While it is possible that a more optimal fluorogenic dye (higher brightness and/or solution K _L–Z_) might improve the utility of construct 290cpHTx, we resolved to continue optimizing the 1–293spHTx:JF_585_ and 10–293spHTx:JF_585_ pairings to take advantage of their greater visibility (see **Figure 2c** representative membrane insets) and larger dynamic ranges.

### Hypotonic Shock Response Occurs Irrespective of Osmolyte Identity

We expect the TRAAKspHTx response to be dependent on membrane tension generated by osmotic pressure, rather than the change in concentration of any particular osmolyte. To confirm this, we repeated our JF_585_ screening experiments with mannitol-, rather than NaCl-, based solutions, such that dilution-effected hypotonic shock would not alter extracellular ionic strength. As expected, control pDisHTx experiments showed no change in JF_585_:EGFP brightness in response to hypotonic shock, regardless of induction stimulus (**Supplementary Figure 3a**; **7**, left). In contrast, both 1–293spHTx and 10–293spHTx (**Supplementary Figure 7**, middle, right) exhibited tension-induced dimming, tempered by Δ(osmolarity): e.g., 1–293spHTx iso_300_–hypo_300_ = 6.1% vs 20.8% in mannitol (–155 mOsm) vs NaCl (–200 mOsm) experiments. Dimming also occurred more slowly in the mannitol than the NaCl hypotonic shock experiments (**Supplementary Figures 3f**–**g**), which we similarly attribute to the smaller stimulus magnitude.

### CAAX Farnesylation Motif is Inessential to Sensor Function

While the CAAX farnesylation motif improved channel trafficking to the plasma membrane in HEK293T cells, it also promoted separation of TRAAKHTxs into raft-like domains made more noticeable by cell swelling (**Supplementary Figure 8**). To verify that this resorting was not responsible for hypotonic shock-induced changes in fluorescence, we removed the C-terminal CAAX tag from the two most responsive split HaloTag sensors. Unlike their CAAX-tagged counterparts, constructs 1–293spHT and 10–293spHT expressed both on the plasma membrane and throughout internal membranes in HEK293T cells (see representative cells in **Figures 3**–**6**). Additionally, cells expressing these constructs appeared better adhered to their poly-L-lysine-coated glass substrate, as evidenced by their many peripheral filopodia,^41^ which densely expressed TRAAKspHTs. Quantification of hypotonic shock-induced changes in total membrane fluorescence in TRAAKspHT-expressing cells, versus plasma membrane fluorescence in the analogous TRAAKspHTx cells, yielded an indistinguishable tension response: 1–293spHT (x) iso_300_–hypo_300_: 21.6 ± 2.9% (20.8 ± 2.1%); 10–293spHT(x) iso_300_–hypo_300_: 20.6 ± 2.5% (16.5 ± 2.0%) mean ± s.e.m. (**Supplementary Figures 9a–b**, **10**). No difference in the relative JF_585_:EGFP dimming of the internal membranes and plasma membrane was observed in TRAAKspHT-expressing cells (**Supplementary Figure 11**), when these membranes could be distinguished.

**Figure 3:**
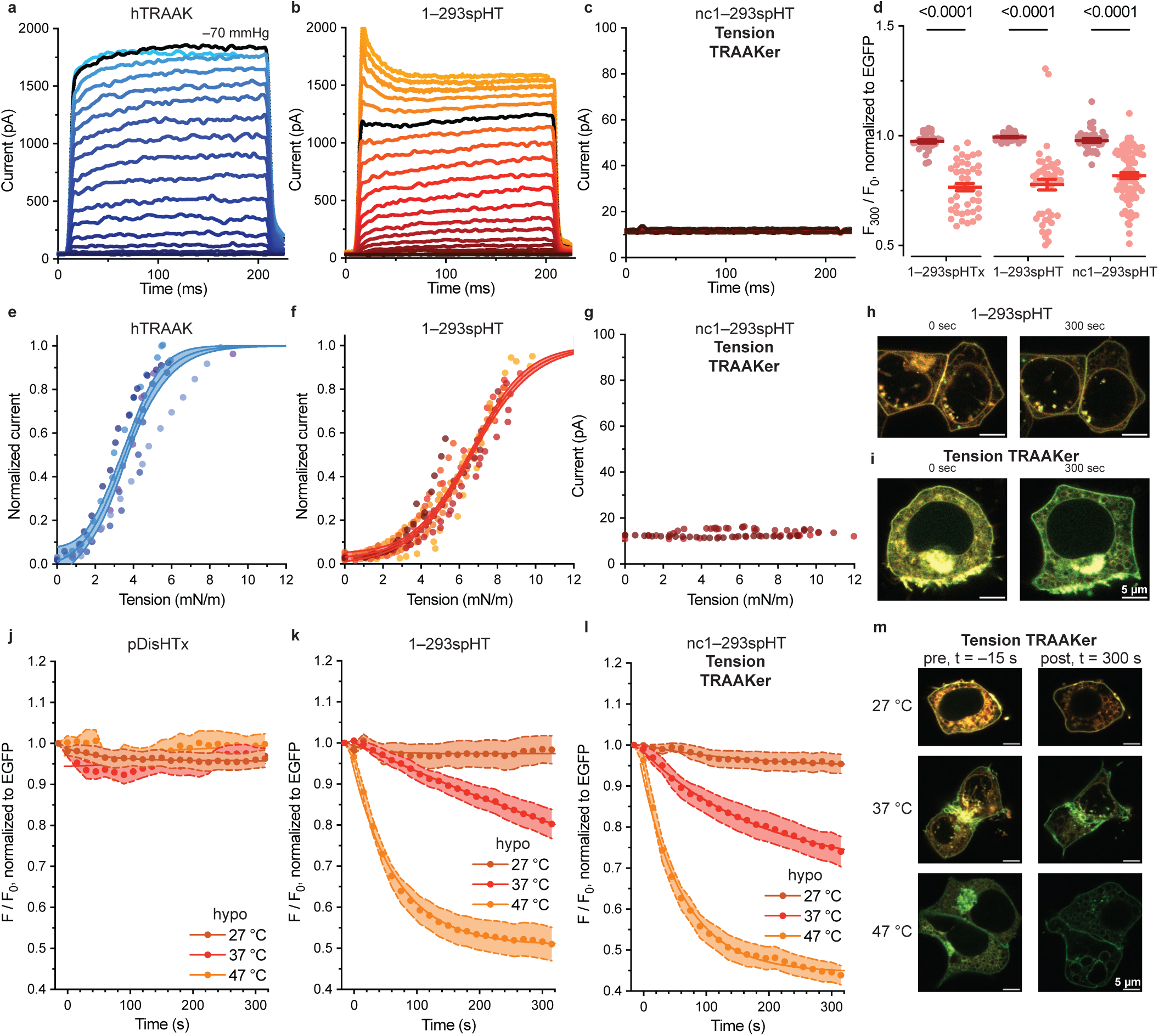
Nonconducting TRAAK spHaloTags respond to hypotonic shock. (**a**–**c**) Representative currents produced by hTRAAK, 1–293spHT, and 10–293spHT, respectively, in excised inside-out HEK293T cell patch in response to increasingly negative pressure steps. Applied pressure steps increase in magnitude in increments of 5 mmHg from dark to light, with currents elicited by a –70 mmHg pressure step highlighted in black. (**d**) Change in brightness of JF_585_ relative to EGFP after 5 minutes in membranes of HEK293T cells transiently transfected with described constructs. Iso/hypo: isotonic/hypotonic solution added. Data shown as mean ± s.e.m. with individual n’s as circles. n ≥ 30. p values calculated from multiple unpaired Mann-Whitney tests. (**e**–**g**) Tension activation curves plotted for constructs hTRAAK (n = 5), 1–293spHT (n = 8), and nc1–293spHT (n = 3), respectively, according to best Boltzmann sigmoidal fit (mean ± s.e.m., solid lines with shaded 95% confidence interval). nc1–293 data is neither normalized nor fit. Data from individual inside-out patches shown as circles, colored by cell. (**h**–**i**) Images of representative cells from hypo condition presented at listed time points for 1–293spHT and nc1–293spHT, respectively. JF_585_ depicted in red, EGFP in green. Brightness scaling maintained within same channel and cell. Distance scale bar = 5 µm. (**j**–**l**) Change in brightness of JF_585_ relative to EGFP over time in membranes of HEK293T cells transiently transfected with pDisHTx, 1–293spHT, or Tension TRAAKer, respectively, at the described temperatures. Hypo: hypotonic solution added <2 seconds prior to t = 0 s (–200 mOsm). Data shown as mean ± s.e.m. n ≥ 7. (**m**) Images of representative Tension TRAAKer cells from hypo condition presented at listed time points and temperatures. Brightness scaling maintained within same channel throughout all cells.

### Split HaloTag Insertion Right Shifts TRAAK’s Tension Activation Curve

The split HaloTag protein augments the size of the human TRAAK channel by approximately 300 residues. We expected this bulky addition would increase the energetic barrier to tension-induced TM4 movement and lateral expansion of the channel, thereby altering TRAAK’s activation curve. To confirm, we characterized the tension–current relationship of both 1–293spHT and 10–293spHT in excised inside-out HEK293T cell patches as previously described.^18^ For comparison, we first examined a human TRAAK–EGFP conjugate, measuring a half-maximal activation (T_50_) of 3.5 ± 0.2 mN/m, with a 10 to 90% activation range of 1.4–5.9 mN/m (**Figure 3a**, **3e**). In contrast, 1–293spHT’s and 10–293spHT’s activation curves are right-shifted to T_50_ = 6.7 ± 0.1 mN/m and 6.5 ± 0.1 mN/m, respectively (**Figure 3b**, **3f**; **Supplementary Figure 12a**–**b**). Boltzmann sigmoidal fits of 1–293spHT and 10–293spHT patch data also reveal that tension-dependent current flux through 1–293spHT occurs with a gentler slope than for TRAAK-EGFP or 10–293spHT (slope factor 1.1 ± 0.1 vs 1.5 ± 0.1 vs 1.1 ± 0.1 for TRAAK-EGFP, 1–293spHT, and 10–293spHT, respectively, **Supplementary Table 2**). As such, 1–293spHT has a broader 10 to 90% activation range (3.0–10.0 mN/m) than 10–293spHT (3.7–8.9 mN/m).

### Nonconducting TRAAKspHTs Report Tension

Potassium conduction through TRAAK is unnecessary, and largely unwanted, in a broadly applicable tension sensor. As such, we endeavored to mutate the channel selectivity filter to ablate ion flux without compromising tension-induced conformational changes. Based on mutational studies of the Shaker potassium channel,^42^ we changed the G(Y/F)G motifs in each TRAAKspHT monomer to GPG (glycine–proline–glycine), hypothesizing that proline insertion would disrupt the potassium coordination environment required for conduction. We verified that neither proline mutant (1–293spHT nor 10–293spHT) produced potassium currents at any tension up to membrane lysis in excised patches (**Figure 3c**, **3g**; **Supplementary Figure 12c**). We termed these nonconducting constructs nc1–293spHT and nc10–293spHT, respectively. Both constructs continued to exhibit tension-induced dimming in JF_585_:EGFP fluorescence in hypotonic shock experiments (**Figure 3d**, **3h**–**i**; **Supplementary Figures 9c–d**, **10**). Like their conducting counterparts, nc1–293spHT and nc10–293spHT were responsive to hypotonic shock regardless of the presence/absence of a C-terminal CAAX farnesylation motif (**Supplementary Figures 9e–f**, **10**). Likewise, hypotonic shock experiments performed with mannitol-based solutions also elicited JF_585_ dimming (**Supplementary Figure 9e**–**f**). In light of the very slightly improved hypotonic shock responses observed for nc1–293spHT compared to nc10–293spHT, as well as 1–293spHT’s broader activation range, we proceeded to characterize the 1–293spHT construct and its nonconducting counterpart, which we hereafter call Tension TRAAKer.

### TRAAKspHT Response is Temperature Dependent

TRAAK is thermosensitive in intact cell recordings, with an open probability that nearly triples between 37 °C and 42 °C.^43^ To characterize Tension TRAAKer’s efficacy at different temperatures, we maintained both cells and solutions at either 27 °C, 37 °C, or 47 °C for at least 10 minutes prior to data collection. Control experiments performed with pDisHTx showed no temperature dependence in JF_585_:EGFP fluorescence, regardless of hypotonic or isotonic solution addition (**Figure 3j**, **Supplementary Figure 13a**). In contrast, 1–293spHT exhibited a significant temperature-dependent increase in response magnitude upon hypotonic shock (**Figure 3k**, **Supplementary Figure 13b**). While cells held at 27°C dimmed by only 1.5% after 5 minutes of exposure to hypotonic solution, this response increased to 18.9% at 37°C, and 48.4% at 47°C. Exponential fit of this data suggests that maximum sensor response to a 200 mOsm osmolarity drop is 49.3% (**Supplementary Table 3**). Tension TRAAKer also exhibited a temperature-dependent response, dimming by 4.5% at 27°C, 24.9% at 37°C, and 55.4% at 47°C (**Figure 3l**), with maximum response predicted to be 55.4% (**Supplementary Table 3**).

In addition to being more responsive to hypotonic shock at higher temperatures, Tension TRAAKer is also basally temperature sensitive in isotonic conditions. Representative cells collected with the same imaging settings and depicted to the same scale in **Figure 3m** demonstrate how JF_585_ fluorescence dims relative to EGFP at higher temperatures. Quantification of this effect shows that while 1–293spHT and Tension TRAAKer exhibit similar initial whole cell JF_585_:EGFP brightness at 27 °C and 37 °C, both dim by almost 70% at 47 °C. In contrast, pDisHTx is slightly brighter in the JF_585_ channel at 47 °C (**Supplementary Figure 14**). This temperature dependence is distinct from the tension response depicted in **Figure 3k** and **3l**, as cells held at temperature for 5 minutes in isotonic solution maintain stable JF_585_:EGFP brightness (**Supplementary Figure 13c**).

### Characterizing Tension TRAAKer:JF_585_

#### Responsiveness, Efficacy vs Flipper-TR®

We next confirmed that Tension TRAAKer’s hypotonic shock-induced dimming is dependent on stimulus magnitude. Holding all cells and solutions at 37°C, we added solution to alter bath osmolarity by either +200, 0, –100, or –200 mOsm, respectively, and monitored resultant changes in JF_585_:EGFP brightness. As in all prior experiments, cells maintained in isotonic solution showed no change in brightness and no noticeable change in shape or behavior. Hypertonicity, in contrast, elicited rapid cell shrinking (**Figure 4b**, **4d**, top) and a small increase in JF_585_:EGFP fluorescence both for 1–293spHT and Tension TRAAKer (**Figure 4a**, **4c**). We speculate this is a result of membrane tension decrease from basal levels upon cell shrinking. This effect was not commensurate to equal and opposite hypotonicity; i.e., hyper_300_–iso_300_ vs iso_300_–hypo_300_ = 8.9% vs 21.8% for 1–293spHT, 7.5% vs 33.4% for Tension TRAAKer. More mild hypotonic shock produced an intermediate effect at 8.8% and 12.9% for 1–293spHT and Tension TRAAKer, respectively.

**Figure 4:**
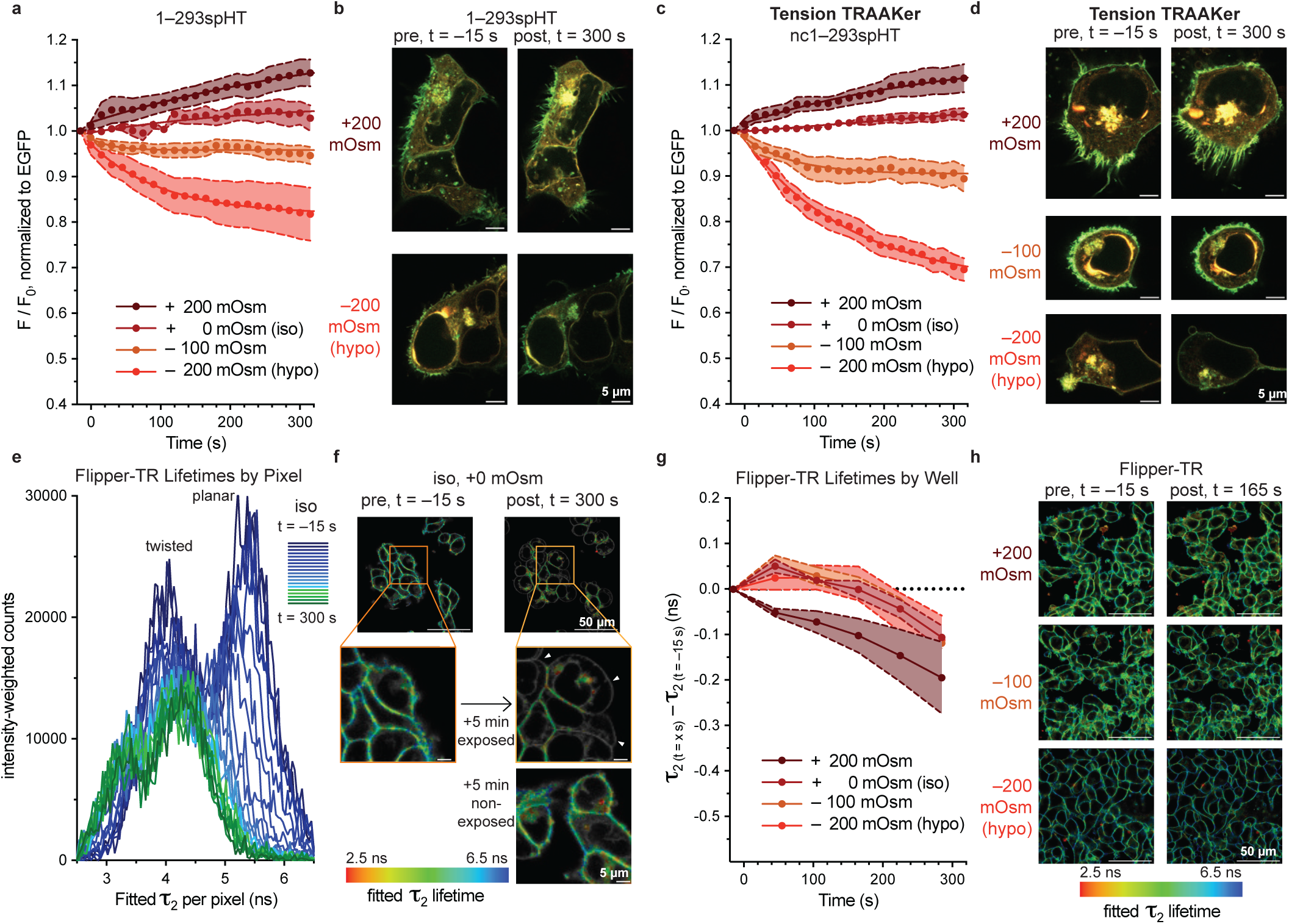
Tension TRAAKer reports tension changes with greater precision than Flipper-TR®. (**a**, **c**) Change in brightness of JF_585_ relative to EGFP over time in membranes of HEK293T cells transiently transfected with 1–293spHT or Tension TRAAKer, respectively, subjected to the listed osmolarity changes <2 seconds prior to t = 0 s. Iso, hypo conditions match previous figures. Data shown as mean ± s.e.m. n ≥ 7 and collected at 37 °C. (**b**) Images of representative 1–293spHT cells from hypertonic (+200 mOsm) and hypotonic (–200 mOsm) conditions presented at listed time points. (**c**) Images of representative Tension TRAAKer-expressing cells subjected to listed conditions presented at listed time points. JF_585_ depicted in red, EGFP in green. Brightness scaling maintained within same channel and cell. Distance scale bar = 5 µm. (e) Intensity-weighted plot of *τ*_2_ lifetimes extracted on a per pixel basis from Flipper-TR®-treated HEK293T cells subjected to isotonic solution addition <2 seconds prior to t = 0 s. Individual pixels were fit to a biexponential model to remove the short lifetime autofluorescent component; *τ*_2_ is the longer lifetime. (f) Images of the experiment described in **e** collected at the listed time points depicted as an overlay of intensity (8-bit black and white image, scaled to 2 photons/bit) and *τ*_2_ (RGB image, color coded as indicated). Representative enlarged images shown at t = –15 seconds and t = 300 seconds both with and without prior light exposure on the field of view. Photo-toxicity is evidenced by cell blebbing in isotonic conditions after repeated imaging (arrowheads). (**g**) Change in effective lifetime of Flipper-TR® over time in wells of HEK293T cells, subjected to the listed osmolarity changes <2 seconds prior to t = 0 s. Iso, hypo conditions match previous figures. Data shown as mean ± s.e.m. n = 3 and collected at 37 °C. Isotonic and hypotonic conditions were statistically indistinguishable in these experiments. (**h**) Images of representative Flipper-TR®-treated HEK293T cell wells subjected to listed conditions presented at listed time points. Images presented as an overlay of intensity (8-bit black and white image, scaled to 2 photons/bit) and *τ*_2_ (RGB image, color coded as described, bin 2). Distance scale bar = 50 µm.

To benchmark Tension TRAAKer’s efficacy compared to available tension sensors, we repeated the identical experiment with HEK 293T cells treated with Flipper-TR®.^23^ Notably, attempts to visualize Flipper-TR® lifetimes at a similar temporal (**Figure 4e**–**f**) resolution to Tension TRAAKer failed due to photobleaching of the probe (**Figure 4e**) and photo-induced injury to the cells (**Figure 4f**). In particular, we noted a disappearance of the longer lifetime component attributed to the planar Flipper-TR® rotamer^23^ from cell membranes while imaging in isotonic conditions, along with the appearance of a short lifetime species in cell interiors (**Figure 4e**). We also observed severe cell blebbing in isotonic conditions within the imaged area, that was absent from neighboring non-exposed cells, consistent with photo-toxicity (**Figure 4f**).

To collect sufficient photons for lifetime estimation while minimizing photo-toxicity, we reduced our effective frame rate to 1/4^th^ of that used for Tension TRAAKer, collecting data for 10 seconds once per minute (compared to 6.45 seconds every 15 seconds for Tension TRAAKer). Under these conditions, we could reasonably fit a double exponential curve to each full field of view of cells (>10^6^ photons), quantifying changes in Flipper-TR® lifetime on a well-by-well basis, rather than cell-by-cell as for Tension TRAAKer (**Figure 4g**). We observed Flipper-TR® lifetimes decreased by ∼0.2 ns in +200 mOsm hypertonic solution (decreased tension) and transiently increased by up to ∼0.06 ns in hypotonic solution (increased tension), consistent with previously reports.^23^ Isotonic and hypotonic conditions were statistically indistinguishable in these experiments. In all conditions we noted photo-induced drift downward in lifetime by the fifth image, along with photo-induced injury to the cells. The spatial resolution of Flipper-TR® tension reporting was also clearly limited relative to Tension TRAAKer. Color-coded images in which Flipper-TR® lifetimes were plotted on a pixel-by-pixel basis required binning to maintain >10^3^ photons per fit (bin = 2, **Figure 4h**), yielding images with 1/150^th^ of the resolution demonstrated for Tension TRAAKer (**Figure 4b**, **d**).

These data demonstrate Tension TRAAKer’s improved dynamic range over Flipper-TR®, particularly to hypotonic shocks, and support its function as a sensor of membrane tension, rather than osmolarity. That is, the tension-conduction relationships described in **Figure 3** suggest that TRAAK is mostly in the TM4 down state at basal/isotonic tensions. While hypertonic shock is expected to reduce membrane tension, the dynamic range encompassing TRAAK closing at low tensions is small. In contrast, hypotonic shock (increased tension) can potentially elicit the TM4 up state in almost all channels from rest, yielding a much larger dynamic range. Flipper-TR®, which probes membrane viscosity, does not exhibit a comparable dynamic range for increased tensions.

#### Reversibility

TRAAK reversibly opens to the mechano-activated open state under tension and rapidly closes (**τ**_close_ ∼ 0.28 ms)^30^ upon cessation of mechanical stimuli. Aggressive cell swelling can induce egregious, unresolvable changes in cell health, but on short time scales is reversible by recovery in isotonic solution.^61^ We performed a series of recovery experiments on Tension TRAAKer and its conducting counterpart, wherein we returned cells to isotonic solution after a period of hypotonic shock while monitoring JF_585_:EGFP fluorescence rebound. **Figure 5a** shows that 1–293spHT JF_585_:EGFP fluorescence remains stable in isotonic solution over 10 minutes but steadily decreases under hypotonic conditions over the same period, with an estimated maximal response of 44% (calculated from exponential decay fit, **Supplementary Table 4**). In contrast, isotonic recovery at 2 minutes or 5 minutes induces JF_585_:EGFP fluorescence recovery (representative cells shown in **Figure 5c**). Whether the transient dimming observed immediately after recovery solution addition is due to increased tension—perhaps due to the rapid inward membrane movement observed upon hypertonic shock—or a consequence of plasma membrane collapse into the field of view is unknown and requires further investigation.

**Figure 5:**
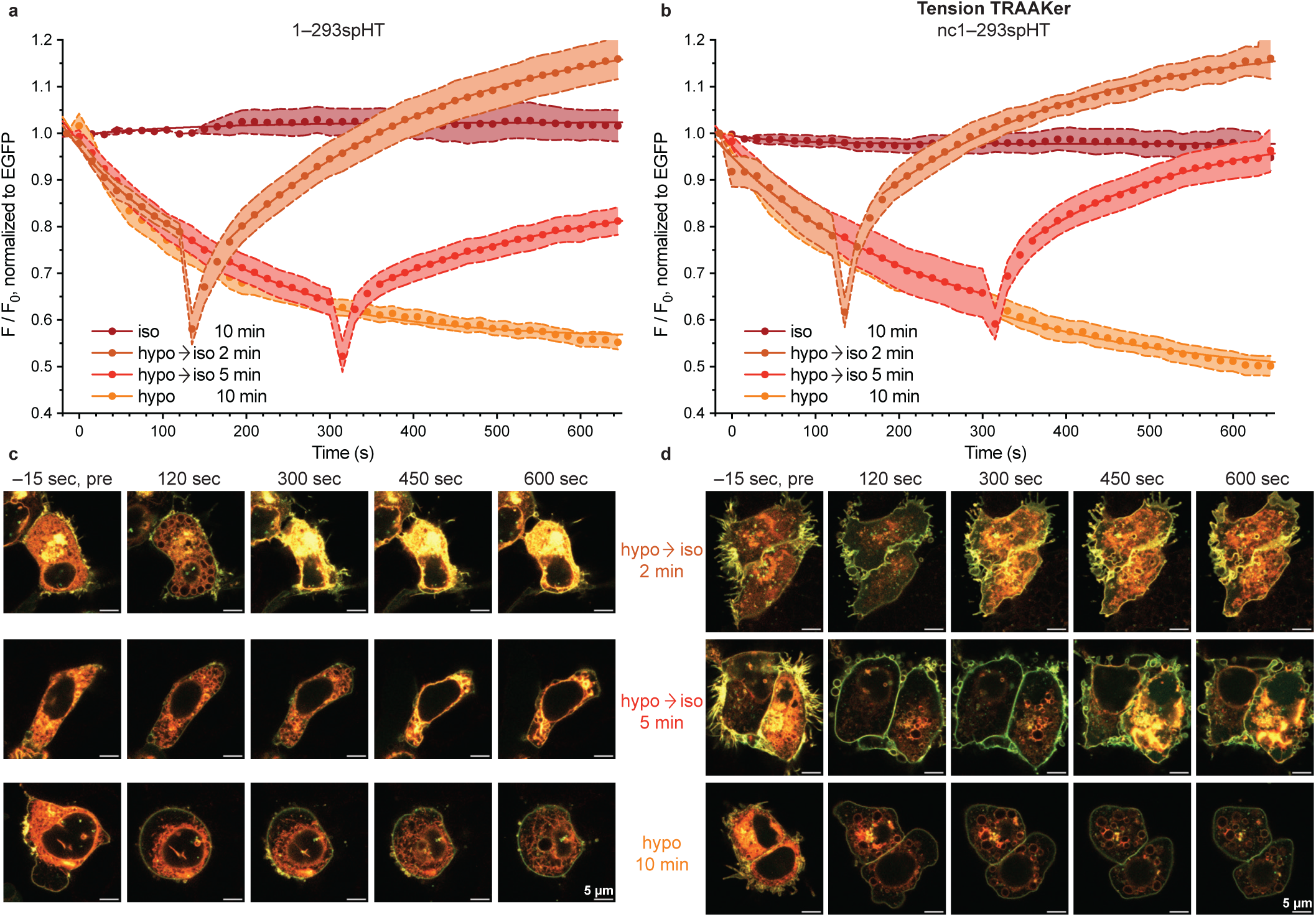
Tension TRAAKer response is reversible. (**a**–**b**) Change in brightness of JF_585_ relative to EGFP over time in membranes of HEK293T cells transiently transfected with 1–293spHT or Tension TRAAKer, respectively, at 37 °C. Iso/hypo: isotonic/hypotonic solution added <2 seconds prior to t = 0 s; data collected for 10 minutes. Hypo → iso: cells returned to isotonicity after hypotonic shock at listed time point. Data shown as mean ± s.e.m. n ≥ 8. (**c**–**d**) Images of representative 1–293spHT and Tension TRAAKer cells, respectively, hypo → iso 2 min, hypo → iso 5 min, hypo 10 min conditions presented at listed time points. JF_585_ depicted in red, EGFP in green. Brightness scaling maintained within same channel and cell; scaling varies from cell to cell. Distance scale bar = 5 µm.

Cells exposed to hypotonic conditions for only 2 minutes recover quickly (∼4.25 minutes), with a subsequent overshoot (exponential fit plateau = 1.24 ± 0.06, **Supplementary Table 5**). After 5 minutes of hypotonic shock, 1–293spHT brightness recovers more slowly, to only 80% of initial after 5 minutes (exponential fit plateau = 0.88 ± 0.13, **Supplementary Table 5**). Tension TRAAKer is similarly responsive to 10 minutes of hypotonic shock, with a calculated maximal response of 53% (**Figure 5b**, **Supplementary Table 4**). It likewise reversibly dims and brightens according to hypotonic shock and recovery, rebounding from 2 minutes under hypotonic conditions within 2 minutes, and from 5 minutes within 5 minutes—both faster than 1–293spHT (**Figure 5b**, **5d**; **Supplementary Table 5**). From these data, we posit that potassium flux through 1–293spHT during cell swelling may interfere with cell recovery, a feature mitigated by the nonconducting Tension TRAAKer. We expect that the over-recovery observed after short hypotonic shocks arises from the increase in membrane surface area promoted by cell swelling, leading to lower overall membrane tensions after isotonic recovery.^44^

#### Spatiotemporal Precision

While membrane tension rapidly equilibrates across simple lipid vesicles, recent studies demonstrate that tension in cellular membranes can be highly spatially and temporally compartmentalized. This localized restriction has profound implications for how cells process distinct mechanical cues^19^. We therefore asked whether Tension TRAAKer could report on tension variation within cellular compartments and membranes.^45,46^ Three-dimensional reconstructions of HEK293T cells expressing Tension TRAAKer pre- and post-hypotonic shock suggest differential tension responses within the plasma membrane and internal compartments (**Supplementary Figure 15**). To more precisely quantify these changes, we imaged cells subjected to hypotonic shock at 2 Hz and examined the resultant JF_585_:EGFP signal per pixel in each frame (**Figure 6a, Supplementary Figure 16**). We then compared different areas of the cell membranes by averaging pixels within a given region of interest. While all membranes progressively dimmed in the JF_585_ channel in response to hypotonic shock, the rate of dimming varied among membrane regions, with bleb formation, for example, preceded by a small local tension increase (**Figure 6b**). These data suggest that Tension TRAAKer reports membrane properties with spatiotemporal precision.

**Figure 6:**
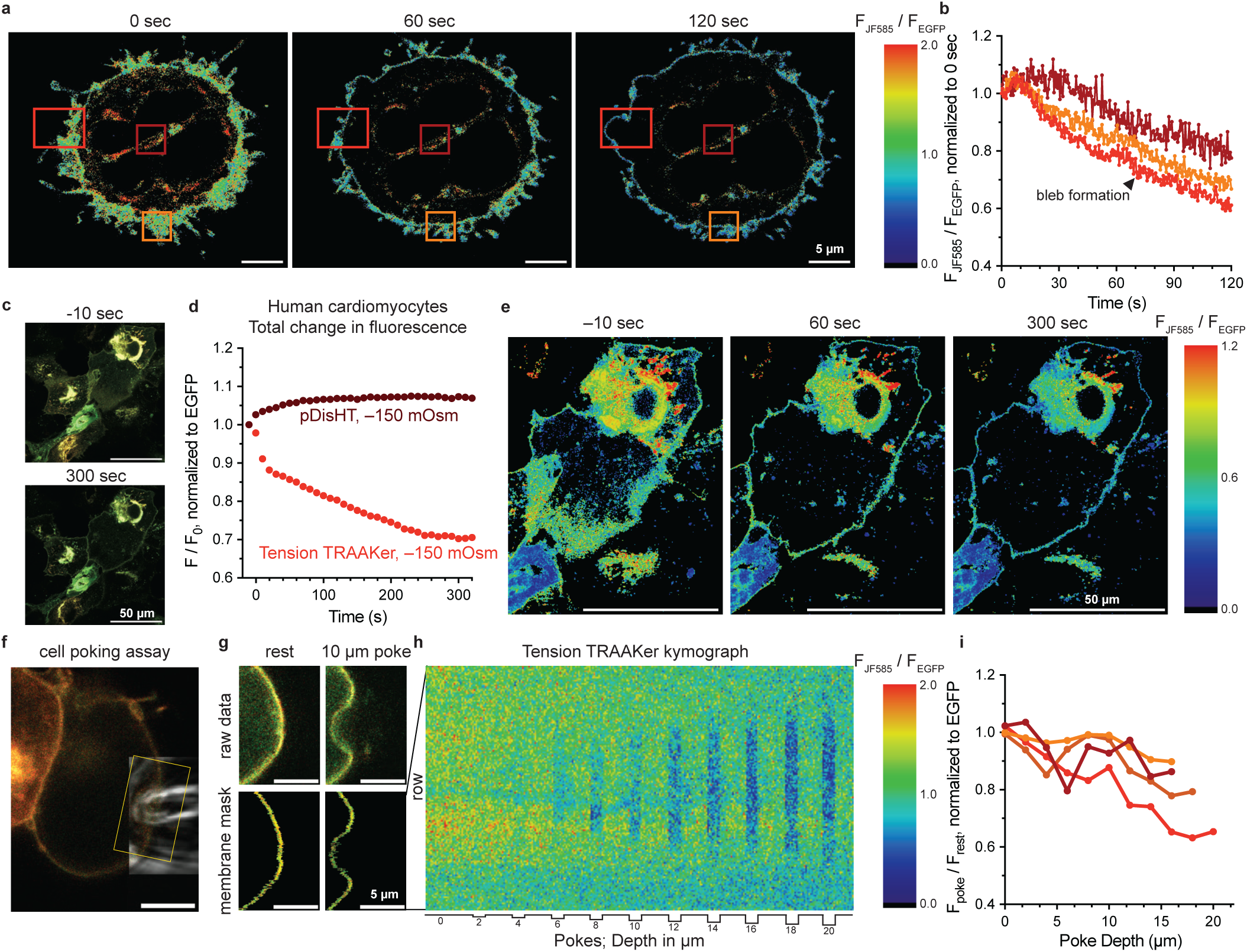
Tension TRAAKer reports tension with high spatiotemporal precision in HEK293T cells and cardiomyocytes. (**a**) Representative Tension TRAAKer-expressing HEK293T cell subjected to –200 mOsm hypotonic shock at 37 °C color coded by JF_585_/EGFP pixel value at listed time points post-hypotonic shock. (**b**) Average change in JF_585_/EGFP pixel value within depicted regions of interest, colored according to boxes in **a**. Bleb formation noted with black arrow. (**c**) Tension TRAAKer-expressing human cardiomyocytes depicted at listed time points before and after hypotonic shock. JF_585_ depicted in red, EGFP in green. Brightness scaling maintained across images. Distance scale bar = 50 µm. (**d**) Change in brightness of JF_585_ relative to EGFP over time in membranes of human cardiomyocytes transiently transfected with Tension TRAAKer or pDisHT (control) at 37 °C after hypotonic shock <2 seconds prior to t = 0 s. (**e**) Representative Tension TRAAKer-expressing human cardiomyocyte subjected to –150 mOsm hypotonic shock at 37 °C color coded by JF_585_/EGFP pixel value at listed time points. (**f**) Representative image of cell poking assay. JF_585_ depicted in red, EGFP in green, glass pipette overlaid in transmitted light. Example ROI shown in yellow box. (**g**) Representative ROI chosen to examine cell poke shown on top; membrane mask of same depicted below. (**h**) Kymograph of a Tension TRAAKer-expressing HEK293T cell subjected to 5-second-long cell pokes of increasing depth from left to right. Each column is a laterally compressed membrane mask of a single frame of video, constructed from the average JF_585_/EGFP value per masked row of pixels (color coded by JF_585_/EGFP value), illustrating the spatiotemporal precision and reversibility of the Tension-TRAAKer response. (**i**) Plot of change in JF_585_/EGFP brightness according to poke depth in Tension TRAAKer-expressing cells (n = 4). JF_585_/EGFP value calculated for entire membrane mask encompassing poke area, normalized to same region prior to poke. Data color coded according to individual cells depicted in **Supplementary Figure 17**.

To validate Tension TRAAKer function in other cell types, we next examined its response to hypotonic shock in human iPSC-derived cardiomyocytes.^47^ As in HEK293T cells, Tension TRAAKer expressed in the plasma membrane of human cardiomyocytes, and prominently in internal membranes, e.g., endoplasmic reticulum (**Figure 6c**). Cardiomyocytes were maintained in JF_585_-containing media at 37°C and subjected to hypotonic shock (approximately –150 mOsm) to increase membrane tension. Tension TRAAKer-expressing cells showed a ∼30% decrease in JF_585_:EGFP fluorescence over 5 minutes (**Figure 6d**), which was absent in the pDisHT-expressing control cells. Analogous to HEK293T cells, Tension TRAAKer dimmed in all cardiomyocyte membranes in response to hypotonic shock. Visualization of the change in JF_585_:EGFP fluorescence per pixel—after down sampling to reduce noise—demonstrated varied responses to hypotonic shock among membrane regions of cardiomyocytes, similar to what we observed in HEK293T cells (**Figure 6e**). These data show that Tension TRAAKer’s response to increased tension is generalizable to other cells.

To probe the limits of Tension TRAAKer’s capacity for monitoring local membrane tension changes, we constructed a cell poking assay wherein a sealed glass pipette is rapidly lowered onto a HEK293T cell by a piezo-driven actuator, held for five seconds, and then raised (**Figure 6f**). In cells with an intact cytoskeleton, cell poking is expected to exert local tension with very limited propagation beyond the stimulus site.^46^ The force exerted is dose dependent; deeper pokes elicit larger TRAAK^48^ and PIEZO1/2^49^ currents in whole-cell patch clamp experiments. To evaluate spatial precision with Tension TRAAKer, we applied cell pokes of increasing depth, climbing from 0 µm in increments of 2 µm, while monitoring fluorescence output. We used the recorded EGFP signal to locate and mask the cell membrane throughout our video recordings (**Supplementary Figures 17a**–**d**) to account for poking-induced membrane deformation (**Figure 6g**). Within this mask, we quantified JF_585_:EGFP brightness per row of masked pixels to condense each video frame into a single column of data.

Chronological alignment of all condensed frames produces a kymograph describing a series of increasing membrane displacements in a given cell (**Figure 6h**, **Supplementary Figures 17e**–**h**). These kymographs show that the magnitude of Tension TRAAKer’s response increases with poke depth, with the largest tensions observed at the stimulus site. In either direction along the membrane proceeding away from the stimulus site—i.e., toward the top or bottom of the kymograph—tension dissipates until Tension TRAAKer’s fluorescent readout is indistinguishable from baseline. Consistent with TRAAK’s previously reported rapid current response to cell poking,^48^ Tension TRAAKer instantaneously dims in the JF_585_ channel upon poke onset, remains dim for the poke duration, and immediately brightens upon membrane relaxation. Quantification of JF_585_ dimming relative to EGFP across the entire region of interest, rather than by row (see ROIs in **Supplementary Figures 17a**–**d**) enables qualitative comparison among different cells (**Supplementary Figures 17i**–**l**), summarized in **Figure 6i**. While evoked tensions are inherently heterogeneous, the poking-elicited local dimming of Tension TRAAKer is universal, with a 10–35% decrease in brightness observed under deep pokes. These data demonstrate that Tension TRAAKer dims in a graded, spatiotemporally precise manner according to indentation depth.

## Discussion

Tension TRAAKer is a membrane protein fluorescent reporter of tension comprising three key genetically encoded components in a single polypeptide chain—the TRAAK transmembrane core, the surrounding split HaloTag, and a C-terminal EGFP—and an applied fluorogenic dye. The TRAAK core is sensitive to a broad range of membrane tensions up to 12 mN/m (i.e., lytic tensions).^27^ Mutation of the selectivity filter to incorporate four proline residues around the channel pore eliminates potassium conduction while maintaining tension sensitivity. According to tension-gated movement of TRAAK’s TM4 helix, the split HaloTag moiety shifts in conformation or position to effect a fluorescence change in the attached environmentally sensitive dye. Inclusion of the C-terminal EGFP allows for precise quantification of this change by ratiometric normalization, irrespective of membrane movement. Together, these modules compose a graded, reversible, spatiotemporally precise fluorescent reporter which responds to multiple membrane tension-inducing stimuli, including osmotic shock and cell poking, in multiple cell types.

Strikingly, both Tension TRAAKer and its conducting counterpart likely report tension changes much smaller than the 10% threshold tension activation value of ∼3.0 mN/m calculated from patch clamp imaging experiments (**Figure 3**). In experiments conducted in HeLa cells, hypoosmotic shocks of ∼200 mOsm corresponded to membrane tension changes of ∼0.2 – 0.8 mN/m, according to optical trapping^50,51^ and Flipper-TR measurements.^23^ Yet, in response to comparable hypotonic shocks in HEK293T cells, Tension TRAAKer dims by up to 50% within 10 minutes. Though limited work has been performed to calculate HEK293T membrane tensions, hypotonic shock-induced formation of migrasomes from retraction fibers (as seen in **Supplementary Figure 15**) requires tension changes of less than 0.1 mN/m.^52,53^ We pose two hypotheses for Tension TRAAKer’s unexpected apparent sensitivity below.

First, it is possible that tension-induced TM4 movement is not the only mechanism for Tension TRAAKer dimming. The C-terminal split HaloTag moiety is connected to both the intracellular end of TM4 and to the remainder of TRAAK’s C-terminal tail, within which a 14-residue Ankyrin G binding sequence^31,32^ connects TRAAK to the cytoskeleton.^54^ Conformational changes induced by hypotonic shock or cell poking might tug on the C-terminal side of the split HaloTag through this cytoskeletal attachment, thereby re-solvating the attached fluorogenic dye and eliciting Tension TRAAKer dimming. We believe this is unlikely due to (1) the 80-residue flexible linker separating the spHT and AnkG motifs and (2) the absence of cytoskeletal elements reported in cell blebs,^55^ within which Tension TRAAKer continues to dim during hypotonic shock experiments (**Figure 6a**–**b**, **Supplementary Figure 16**).

Second, the conformational changes involved in upward movement of a single TRAAK TM4 helix may precede those of both TM4s. TRAAK’s mechano-activated conductive state is characterized by the shifting of both TM4s upward and outward.^28^ However, the movement of only one TM4 helix would be expected to alter spHT:JF_585_ fluorescence. As such, the tension-response curve describing JF_585_ dimming might be left-shifted relative to 1–293spHT’s tension-conduction curve (one TM4 movement vs two). This hypothesis is supported by the observations of both TRAAK^27^ and TREK-1^56^ in one up/one down TM4 conformations by crystallography/cryo-EM. The barrier to movement of one TM4 (which effects spHT:JF_585_ fluorescence change) is seemingly lower than the barrier to the movement of both (which effects potassium conduction).

Tension TRAAKer compares favorably against the only commercially available, albeit indirect, tension reporter Flipper-TR®. While Flipper-TR® is highly sensitive to increases in lipid packing—exhibiting up to a ∼0.5 ns lifetime shift per ∼0.3 mN/m decrease in membrane tension^57^—sensitivity depends strongly on cell identity and membrane composition and saturates above ∼0.5 mN/m (<300 mOsm).^23^ Tension TRAAKer exhibits a significantly larger response at high tensions than Flipper-TR®— Δ33% vs Δ0.03 ns in our hands at 37°C, –200 mOsm. Tension TRAAKer is also able to distinguish among graded hypotonic shocks—Δ12.9 ± 2.6% vs Δ33.4 ± 2.7% at –100 and –200 mOsm, respectively—that are statistically indistinguishable with Flipper-TR® (**Figure 4**). Additionally, Tension TRAAKer can be used to monitor changes in membrane tension with a spatiotemporal precision that is simply impossible with Flipper-TR®, which is practically restricted in resolution by the high photon budget required to accurately calculate fluorescence lifetime.^58,59^

Tension TRAAKer also compares favorably to published noncommercial tension reporters. Chemical sensors like ZNH-Vis^4^ and XZTU-Mito-Vis^60^ have similar limitations to Flipper-TR® in that they respond to changes in membrane viscosity, rather than tension. Mechanosensitive channel-derived tools (e.g., MscL(G22S)61-cpGFP^61^ or GenEPi—a PIEZO1-GCaMP conjugate^62^) have narrower tension responses than Tension TRAAKer, with MscL’s and PIEZO1’s 10 to 90% activation ranges spanning only ∼3 mN/m,^18^ in contrast to 1–293spHT’s 7 mN/m. Opening of both MscL and PIEZO1 also significantly perturbs cell physiology—PIEZO1 is a nonspecific cation channel with a large (∼500 nm^2^) footprint that bends membranes and MscL has ∼40 Å diameter pore permeable to molecules up to 1 kDa in size. In contrast, Tension TRAAKer is nonconductive. Neither MscL(G22S)61-cpGFP nor GenEPi internally control for membrane movement; Tension TRAAKer, on the other hand, has a C-terminal EGFP to precisely quantify channel localization. Moreover, neither GenEPI nor MscL(G22S)61-cpGFP have been used to characterize membrane tension changes following their initial descriptions.

Looking forward, Tension TRAAKer is readily amenable to adeno-associated virus packaging (at 969 residues in length) and *in vivo* transduction, rendering it genetically targetable. JF_585_ has a favorable two-photon cross section (1100 nm), is blood–brain barrier-permeable, enables long-lasting HaloTag visualization, and requires no wash-out (labelling of HaloTag-expressing cortical neurons in mice has been demonstrated within 5 minutes of intravenous tail injection, lasting for nearly 2 weeks).^39^ JF_585_ is also commercially available, as are a number of related fluorogenic dyes. While we found the turn-on properties, photostability, and environmental sensitivity of JF_585_ to be optimal for our work, we expect other low K_L-Z_ fluorogenic dyes will also enable tension monitoring with Tension TRAAKer. Due to its sensitivity, reversibility, and spatiotemporal precision, Tension TRAAKer is a broadly useful addition to the toolbox of fluorescent reporters for the study of membrane tension.

## Methods

### Compounds

All chemical compounds were obtained commercially unless otherwise noted. Janelia Fluor® 635 chloroalkane and Janelia Fluor® 585 chloroalkane were gifted by the Lavis group at Janelia Research Campus. BF_646_ was synthesized by the Miller group at the University of California, Berkeley.^38^ All dyes were diluted in DMSO to make high concentration stocks (≥1 mM), aliquoted in Eppendorf tubes, and frozen at –20°C to reduce freeze-thaw cycles.

### Plasmids

Full length *Homo sapiens* TRAAK (UniProt Q9NYG8-2) was obtained from a previously described eukaryotic expression codon optimized construct^18^ that was conjugated to a C-terminal EGFP via thrombin cut site and inserted into a pNEH backbone with a CMV mammalian expression promoter. This backbone was used for all CAAX-bearing constructs. pDisplay, HaloTag, and split HaloTag were cloned from HaloTag–pDisplay IRES mCherry.^63^ Circularly permuted HaloTag was cloned from pCAG-HASAP, which was a gift from Eric Schreiter (Addgene plasmid #138325; http://n2t.net/addgene:138325; RRID:Addgene_138325).^25^ The CAAX farnesylation motif was cloned from EGFP-CAAX, which was a gift from Lei Lu (Addgene plasmid #86056; http://n2t.net/addgene:86056; RRID:Addgene_86056).^33^ For all CAAX-free constructs, the backbone from mOrange-N1 was used (identical to pNEH, employed for ease of restriction digest), which was a gift from Michael Davidson (Addgene plasmid #54499; http://n2t.net/addgene:54499; RRID:Addgene_54499).^64^ Full sequences of all constructs discussed herein appear in the Supplementary Information. Nonconducting TRAAK channels were made via sequential site-directed mutagenesis using the following primers:

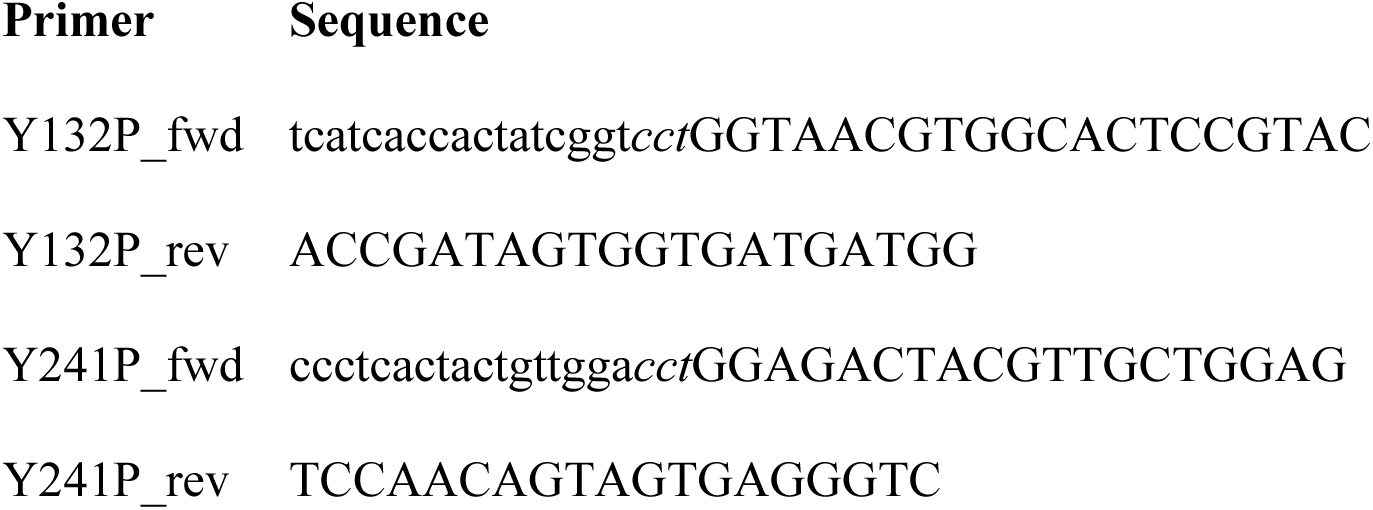

### Human Embryonic Kidney (HEK) 293T Cell Culture and Transfection

HEK293T cells were purchased from the University of California, Berkeley Cell Culture Facility (RRID SCR_017924). Cells were grown on 10 cm tissue culture dishes in DMEM, high glucose (Fisher Scientific) supplemented with 10% fetal bovine serum (Invitrogen), 100 U/mL penicillin-streptomycin (Fisher Scientific), and 1X MEM Non-Essential Amino Acids Solution (Fisher Scientific). Cells were maintained in a 37 °C, 5% carbon dioxide, 96% relative humidity incubator and passaged every three to four days. Passaging of cells was accomplished by aspiration of media, washing with Dulbecco’s phosphate-buffered saline, treatment with 2 mL of 0.25% trypsin-EDTA (Fisher Scientific) until full dissociation of cells was observed (approximately two minutes), and dilution with 8 mL of growth medium. Cells were routinely passaged at 1 in 20 to 1 in 10 dilution.

For confocal imaging experiments, HEK293T cells were grown to >50% confluency on 6 well tissue culture dishes in 2 mL growth medium. Transfection was performed according to manufacturer’s instructions with 1 µg of construct and 3 µL of FuGENE® 6 (Promega) in 45 µL Opti-MEM^TM^ (Invitrogen). Transfected cells were passaged once—as described above, at ∼80–95% confluency—between transfection and imaging to transfer and dilute construct-expressing cells onto poly-L-lysine-treated (0.1% (w/v) in H_2_O, Millipore Sigma) 18 well µ-slides on #1.5 glass (Ibidi). Confocal imaging experiments were performed 40–56 hours post-transfection to maximize construct expression.

For patch clamp electrophysiology experiments, HEK293T cells were seeded directly onto poly-L-lysine-treated (0.1% (w/v) in H_2_O) 12 mm circular glass coverslips (Fisher Scientific) in 6 well tissue culture dishes at low confluency prior to transfection. After re-adherence (>6 hrs), cells were similarly transfected with FuGENE® 6 and 1 µg of construct according to manufacturer’s instructions. Wild-type TRAAK–EGFP was co-transfected with EGFP-CAAX to improve membrane visibility; TRAAK spHT constructs were sufficiently bright to visualize absent EGFP-CAAX co-expression. Excised patch clamp electrophysiology experiments were performed 18–36 hours post-transfection to minimize cell confluency. For cell poking experiments, HEK293T cells were grown to >50% confluency on 6 well tissue culture dishes in 2 mL growth medium. Transfection was performed according to manufacturer’s instructions. Transfected cells were passaged once—as described above, at ∼80–95% confluency—between transfection and imaging to transfer and dilute construct-expressing cells onto poly-L-lysine-treated (0.1% (w/v) in H_2_O) 35 mm dishes with #1.5 glass bottoms (Mattek). Cell poking experiments were performed 40–56 hours post-transfection to maximize construct expression.

### iCell Human Cardiomyocyte Cell Culture and Transfection

iCell cardiomyocytes were purchased from FujiFilm Cellular Dynamics, Inc. (01434), thawed, and plated at 40,000 cells per well in gelatin-treated (0.1% in water, STEMCELL Technologies) 18 well µ-slides on #1.5 glass (Ibidi) according to manufacturer’s instructions. Cells were maintained in a 37 °C, 5% carbon dioxide, 96% relative humidity incubator and fed every two to three days via the complete replacement of existing with fresh cardiomyocyte media (FujiFilm Cellular Dynamics, Inc.).

For confocal imaging experiments, iCell cardiomyocytes were transfected according to manufacturer’s instructions with 0.2 µg of construct and 0.4 µL of ViaFect^TM^ (Promega) in 20 µL Opti-MEM^TM^ (Invitrogen). Confocal imaging experiments were performed 40–56 hours post-transfection to maximize construct expression and trafficking to the cell membrane.

### Confocal Imaging Screening Experiments

The following solutions were used for these experiments:

**Isotonic**: 135 mM NaCl, 15 mM KCl, 3 mM MgCl_2_, 1 mM CaCl_2_, 10 mM HEPES; pH 7.3 in NaOH

**Hypotonic**: 15 mM KCl, 3 mM MgCl_2_, 1 mM CaCl_2_, 10 mM HEPES; pH 7.3 in NaOH

**Mid-hypotonic**: 1 : 1 isotonic : hypotonic solution

**Hypertonic**: isotonic solution + 150 mM NaCl

**NaCl Stock**: 4.76 M NaCl in hypotonic solution

**Hypotonic, Mannitol**: 25 mM NaCl, 15 mM KCl, 3 mM MgCl_2_, 1 mM CaCl_2_, 10 mM HEPES; pH 7.3 in NaOH

**Isotonic, Mannitol**: hypotonic, mannitol solution + 200 mM mannitol

Osmolarities were measured with a VAPRO® Vapor Pressure Osmometer (ELITechGroup), which was confirmed to be within calibration before each use. Solutions were aliquoted and maintained at room temperature (20–25 °C) unless otherwise noted. All hypo/iso/hypertonic shock experiments were performed with the addition of 3 volumes (3V) of test solution to 1 volume (1V) of isotonic solution.

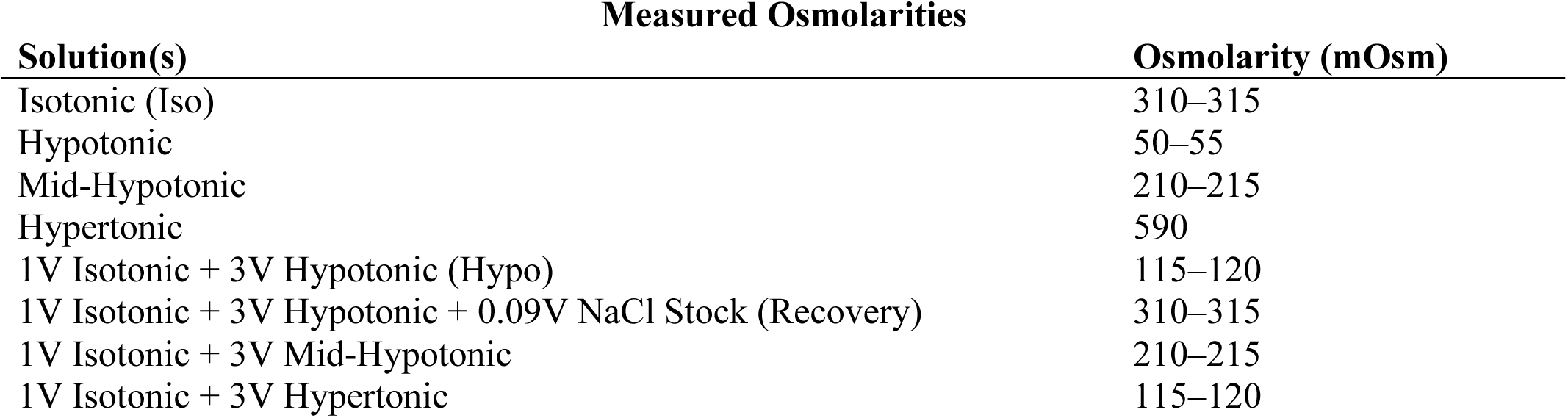
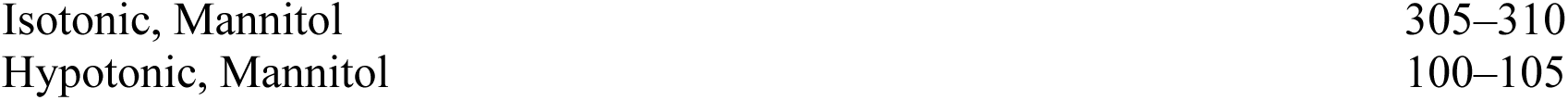

HEK293T cells transfected with construct were treated with 0.5 µM dye in 50 µL of isotonic solution (following a 100 µL isotonic solution rinse) for at least one hour prior to imaging. Cells were imaged on a LSM990 confocal microscope in UC Berkeley’s Molecular Imaging Center. A light blocking incubation chamber maintained at 37 °C was used to maintain cell health and minimize photobleaching. A maximum of two 18 well µ-slides were examined per imaging session during initial screening experiments. All data was collected using a 63x/1.4 PlanApo oil objective to maximize plasma membrane resolution.

Isotonic and hypotonic conditions were achieved via pipette addition of three volumes of the related solution (150 µL) immediately (<5 s) prior to experiment start. Collection of a 4x averaged, 6.45 s/frame, 16-bit, 135 x 135 µm, 1584 x 1584 px, 4-slice z-stack (10 µm in height) every 30 seconds yielded 11 total frames (5 minutes) of data describing construct response (EGFP and dye channels imaged simultaneously with unidirectional scanning). Lasers (488 nm and 561 or 639 nm according to dye) were maintained at the same power settings, but data collection occurred across many days. To account for potential variability in laser power, control isotonic experiments with each construct were performed every day hypotonic/hypertonic test data was collected.

All data was worked up via a custom python script (membrane workup). Briefly, for CAAX-bearing constructs, individual fluorescent cells and indistinguishable cell patches were cropped to separate files in FIJI (ImageJ) via deletion/omission of the unwanted EGFP signal; then masked via application of a blur, threshold, clean, close, dilate, erode, and outer contour sequence to the remaining EGFP pixels. Within the resultant 20-pixel wide membrane mask, the brightest 6 adjacent pixels (“pixel patch”) were chosen along each of a series of lines plotted radially from the inner to outer mask contour to produce an approximately 0.5 µm wide cell membrane. The final thresholded membrane data used in all TRAAK(cp/sp)HTx figures/calculations comprises only pixel patches of average EGFP signal greater than 1500 AU (16-bit scale, approximately 10x background). Every frame of every file was reproduced with only these thresholded points in both the EGFP and fluorogenic dye channels. If the python script omitted a frame of data, or if the produced frame lacked fidelity to the cell membrane (checked manually), the frame was omitted. This most often occurred in instances of cell swelling wherein patches of construct-expressing membrane moved so far apart as to become difficult to mask as a single unit. Data for each cell/patch of cells is reported within this thresholded membrane as:

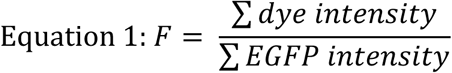

and normalized to t = 0 or t = –15 seconds as described.

While this same python script was initially applied to examine the CAAX-free TRAAKspHTs, the multitude of fluorescent internal membranes within HEK293T cells expressing these constructs rendered isolation of the plasma membrane impossible in most cells. As such, we modified our python script (cell workup) to still mask the EGFP signal via application of a blur, threshold, clean, close, dilate, and erode sequence; but then simply chose all pixels with EGFP signal greater than 1500 AU within the EGFP mask. Data for each cell/patch of cells was reported within the final thresholded mask according to equation 1 above. We estimated internal membrane brightness as:

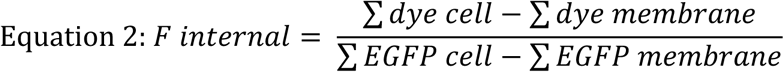

While slight differences in our python scripts render this internal data imperfect, we found it to be a useful approximation of tension changes in internal vs plasma membranes at low time resolution.

A similar, threshold-less script (dye workup) was applied to quantify dye fluorescence relative to background in CAAX-tagged constructs. For single cells with minimal internal construct expression (approximated from EGFP localization), average dye fluorescence per pixel in the membrane (F_membrane_) was calculated using the membrane workup script, while average background fluorescence per pixel (F_background_) was calculated as:

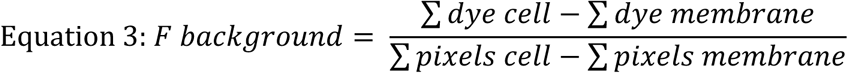

It should be noted that this calculation likely underestimates F_membrane_ / F_background_ due to both the tightness of the membrane mask and the presence of sparse internal channel expression, particularly for spHTx constructs.

Number of distinct slices of distinct (patches of) cells (n) appears for every condition in the source data, as does the number of cells present in each patch (mode = 1).

### Patch Clamp Electrophysiology

Data were measured using the patch-clamp technique with an Axon Axopatch 200B amplifier (Molecular Devices, San Jose, CA). The output of the patch-clamp amplifier was filtered with a built-in lowpass, four-pole Bessel filter having a cutoff frequency of 1 kHz and sampled at 10 kHz. Pulse stimulation and data acquisition used Molecular Devices Digidata 1550B controlled with pCLAMP software version 11 (Molecular Devices, San Jose, CA). Borosilicate glass micropipettes (Sutter Instruments, Novato, CA, internal diameter = 1.2 mm) were fire-polished to a tip diameter yielding a resistance of 1–3 MΩ in the working solutions. The pipette solution was composed of 150 mM KCl, 3 mM MgCl_2_, 5 mM EGTA, and 10 mM HEPES (pH 7.2 with KOH); the bath solution comprised 135 mM NaCl, 15 mM KCl, 3 mM MgCl_2_, 1 mM CaCl_2_, 10 mM HEPES (pH 7.3 with NaOH). Data were collected from n ≥ 3 cells in each condition across several days.

Pressure was applied in incrementally (2–5 mmHg) increasing steps over 200–800 ms with a high-speed pressure clamp device (HSPC-2-SB, ALA Scientific Instruments) while excised inside-out patches were held at 0 mV by the Axopatch 200B. Excised patches were visualized with application of cyan (to excite EGFP) or green (to excite JF_585_) light from an LED light engine (SpectraX, Lumencor), passed through a filter cube and water immersion objective lens (60x, NA1.0).

Videos were recorded at 20–50 Hz with an infrared camera (IR-2000, DAGE-MTI), rotated and cropped in FIJI (ImageJ) to vertically align the membrane, and analyzed via custom python scripts (patch workup) similar to those previously described.^18^ Briefly, the brightest 12 pixels were identified in each row of pixels to mask the cell membrane. Within each row’s 12-pixel mask, the 6^th^ pixel was chosen as the membrane midpoint in that row. When membranes were less than fully brighter than background (i.e., brighter in every row across every video frame), a subset of brightest rows was chosen for workup to eliminate noise. A circle was fit to the resultant midpoint pixels using the Python package circle-fit [https://pypi.org/project/circle-fit], and a patch radius was recorded for each frame of video. Patch radii (r) collected in the middle of each pressure step (at maximal membrane displacement) were averaged, absent the rare outlier resulting from an improperly masked membrane, as checked visually across all frames for all cells. Average radii were used to calculate membrane tension (T) resulting from pressure (P) application using Laplace’s law:

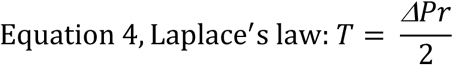

For each excised patch, the average equilibrated current recorded within each pressure step was plotted versus calculated membrane tension and analyzed with Prism 10 (GraphPad Software, LLC).

A Boltzmann sigmoidal curve was fit to each patch with no constraints, but differential weighting—either to 1/current^2^ or unweighted—according to best fit, so as to predict the top of the curve/saturating current. This value was used to normalize the recorded average currents for each patch, as TRAAKspHT patches did not routinely saturation. A plot of normalized current versus tension for all patches of each construct was then fit to a final unweighted Boltzmann sigmoidal curve constrained to top = 1.0 to describe tension-induced channel opening. Number of distinct patches (n) recorded per construct: n = 5, TRAAK; 8, 1–293spHT; 4, 10–293spHT; 3, nc1–293spHT; 3, nc10–293spHT.

### Tension TRAAKer Confocal Characterization Experiments

To precisely account for the temperature and time dependence of Tension TRAAKer responsiveness, all confocal imaging experiments related to 1–293spHT and Tension TRAAKer characterization (and the associated pDisHTx negative control) were conducted with the following alterations: (1) the LSM990 full dark incubation chamber was maintained at temperature with cells inside for at least 10 minutes prior to experiment start, with temperature continuously monitored by a SwitchBot Thermometer Hygrometer (SwitchBot), (2) all solutions were maintained at temperature for at least 10 minutes prior to experiment start, (3) only a single plane of HEK293T cells were imaged in each experiment, (4) images were collected every 15 seconds for up to 10 minutes, with JF_585_:EGFP fluorescence normalized to the image taken at t = –15 seconds, and (5) solution addition was made <2 seconds prior to timepoint 0. All videos were collected using 4x averaging, at 6.45 s/frame, 16-bit, 135 x 135 µm, 1584 x 1584 px. All data was worked up according to the cell workup script described above, including that collected for the pDisHTx negative control. Isotonic recovery, when performed, was initiated by manual addition of 4.4 µL of NaCl stock solution immediately following the listed time point.

For experiments examining the temporal precision of Tension TRAAKer’s response to hypotonic shock, the field of view was shrunk to 34 x 34 µm and videos were collected at 2 Hz with 1x averaging and bidirectional scanning. The timing of solution addition was determined in post by visual artifact within the video. All data was worked up according to the cell workup script. Masked frames were then processed through FIJI’s image calculator, wherein the JF_585_ signal was divided by the EGFP signal, smoothed, and colored according to JF_585_:EGFP signal in each pixel.

### FLIM Experiments with Flipper-TR®-treated HEK293T cells

Fluorescence lifetime imaging (FLIM) experiments were performed on the same LSM990 microscope as all other confocal experiments, with the addition of a Becker-Hickl Time-Correlated Single Photon Counting (TPSC) FLIM system equipped with a pulsed 50 MHz 488 nm excitation laser and 575/150 nm bandpass emission filter. Photons were collected for 10 seconds per image through the same 63x/1.4 PlanApo oil objective to maximize plasma membrane resolution across a 135 x 135 µm, 256 x 256 px field of view. Cells were treated with a 1:1000 dilution of Flipper-TR® (Spirochrome) in 50 µL isotonic solution for at least 15 minutes at 37°C prior to experiment start, according to manufacturer’s instructions. Isotonic, hypotonic, and hypertonic conditions were achieved via pipette addition of three volumes of the related solution (150 µL) equilibrated to 37°C immediately (<2 s) prior to timepoint 0. SPCImage software (Becker & Hickl GmbH) was used to fit fluorescence decay data to a biexponential model in which the longer lifetime (*τ*_2_) was extracted to quantify Flipper-TR® conformation, as previously described.^23^

### Tension TRAAKer Characterization Experiments in iCell Cardiomyocytes

For experiments characterizing the precision of Tension TRAAKer’s response to hypotonic shock in iCell cardiomyocytes, videos were collected across a 135 x 135 µm, 1584 x 1584 px field of view with 1x averaging. Cells were treated with 0.5 µM JF_585_ for at least three hours in iCell cardiomyocyte media (FujiFilm Cellular Dynamics, Inc., 290–320 mOsm) before imaging. Hypotonic shock was effected via the addition of one volume of water to the cardiomyocytes in JF_585_-containing media. As transfected cardiomyocytes universally had many near neighbors, for bulk quantification, images were masked based on a 1500 AU threshold in the EGFP channel and JF_585_:EGFP fluorescence was quantified on an image-by-image basis. For subcellular analysis, images were down sampled by approximately 10-fold to 512 x 512 px, masked to a 5000 AU threshold in the EGFP channel, and then processed through FIJI’s image calculator, wherein the JF_585_ signal was divided by the EGFP signal and colored according to JF_585_:EGFP ratio in each pixel. The higher threshold mask ensured enabled the elimination of non-plasma membrane, non-endoplasmic reticulum fluorescence.

### Cell Poking Experiments

Glass probes for cell poking experiments were made from borosilicate glass micropipettes (Sutter Instruments, Novato, CA) fire polished to seal. The probe was mounted to a piezo-driven actuator driven by a controller/amplifier (P-601/E-625; Physik Instrumente) controlled through a 33220A 20 MHz Function/Arbitrary Waveform Generator (Agilent). The controller was clasped to a MPC-200 micromanipulator (Sutter Instrument) mounted on the LSM990 confocal microscope stage with the full dark incubation chamber maintained at 37 °C. Transfected cells were treated with 0.5 µM JF_585_ in 100 µL of isotonic solution (applied directly to the central glass coverslip after a 300 µL rinse with isotonic solution) for at least one hour prior to imaging, at which point they were diluted in several more milliliter of isotonic solution to achieve sufficient bath height and brought into focus using the 63x/1.4 PlanApo oil objective. The probe was positioned above the side of the cell such that the first, shallowest, downward poke did not produce indentation (0 µm). Each poke was applied for 5 seconds, with probe displacement increasing in increments of 2 µm. Each poke was manually triggered 4–7 frames into a 25-frame, 0.65 s frame rate, 4x averaged, 16-bit, 45 x 45 µm continuous video of the cell membrane (transmission, EGFP, and JF_585_ channels imaged simultaneously with bidirectional scanning).

Videos were rotated and cropped in FIJI (ImageJ) to vertically align the membrane and analyzed via custom python scripts (poke workup) similar to those for the excised patch tension workup. Briefly, in the EGFP channel, the brightest 6 pixels were identified in each row of pixels to mask the cell membrane. Every frame of every video was reproduced with only these 6 pixels in each row in both the EGFP and JF_585_ channels and checked for fidelity to the plasma membrane. This initial screening eliminated all cells with (1) bright internal membranes that extended to the poke site or (2) dim plasma membranes that disappeared from the field of view upon poking. The remaining cells were quantified in two ways. First, sum(JF_585_)/sum(EGFP) was calculated for each row of 6 pixels, enabling lateral compression of each video frame to a single column of data. Plotting of each compressed 25-frame poke video from left to right according to increasing poke depth yielded kymographs describing Tension TRAAKer dimming according to indentation. Second, sum(JF_585_)/sum(EGFP) was calculated for a region of interest comprising the membrane affected by the deepest indentation applied to each cell. Plotting of this data enabled the examination of average Tension TRAAKer dimming across the entire poke site according to indentation onset, and recovery upon offset. To evaluate the dose dependence of Tension TRAAKer dimming according to indentation depth, 5 frames of poke data were averaged and normalized to the average of the preceding 5 frames of rest data (where possible, onset/offset frames excluded).

## Data Availability

Source data are provided with this manuscript. Additional data is available from the corresponding authors upon request.

## Code Availability

Code used to quantify plasma membrane fluorescence, total membrane fluorescence, and membrane tension is available here: https://github.com/BrohawnLab.

## Acknowledgements

This work was supported by National Institutes of Health grants EY035182 (A.V.E.); GM153237 (E.W.M.), NS121431, R01EY024334, and P30EY003176 (R.H.K.); and GM145869 and GM123496 (S.G.B.). A.V.E. was also partially supported by a Weill Neurohub Research Award. Fluorogenic dyes JF_585_ and JF_635_ were provided by Luke D. Lavis at Janelia Research Campus, for which we are immensely grateful. HEK 293T cells stocks were maintained and distributed by the UCB Cell Culture Facility, RRID: SCR_017924. Confocal imaging experiments were conducted at the CRL Molecular Imaging Center, RRID:SCR_017582, funded by NIH S10OD025063. We would additionally like to thank Trevor Docter, for his help with python scripting, and Holly Aaron, Luis Alvarez, Cherise Stanley, and Feather Ives for their imaging support.

## Author Contributions

A.V.E., R.H.K., and S.G.B. conceived the project. A.V.E. and S.G.B. designed TRAAK HaloTag constructs.

A.V.E. performed all experiments and analyzed all data. N.G. and E.W.M. designed and synthesized BF_646_.

B.E.S. established the cell poking apparatus for imaging experiments. A.V.E. and S.G.B. wrote the manuscript with input from all authors.

