## Supplementary Information for "Tension TRAAKer: a chemigenetic fluorescent membrane tension reporter"

### Supplementary Data

1. Supplementary Figures
2. Supplementary Tables
3. Full Construct Sequences

#### 1. Supplementary Figures

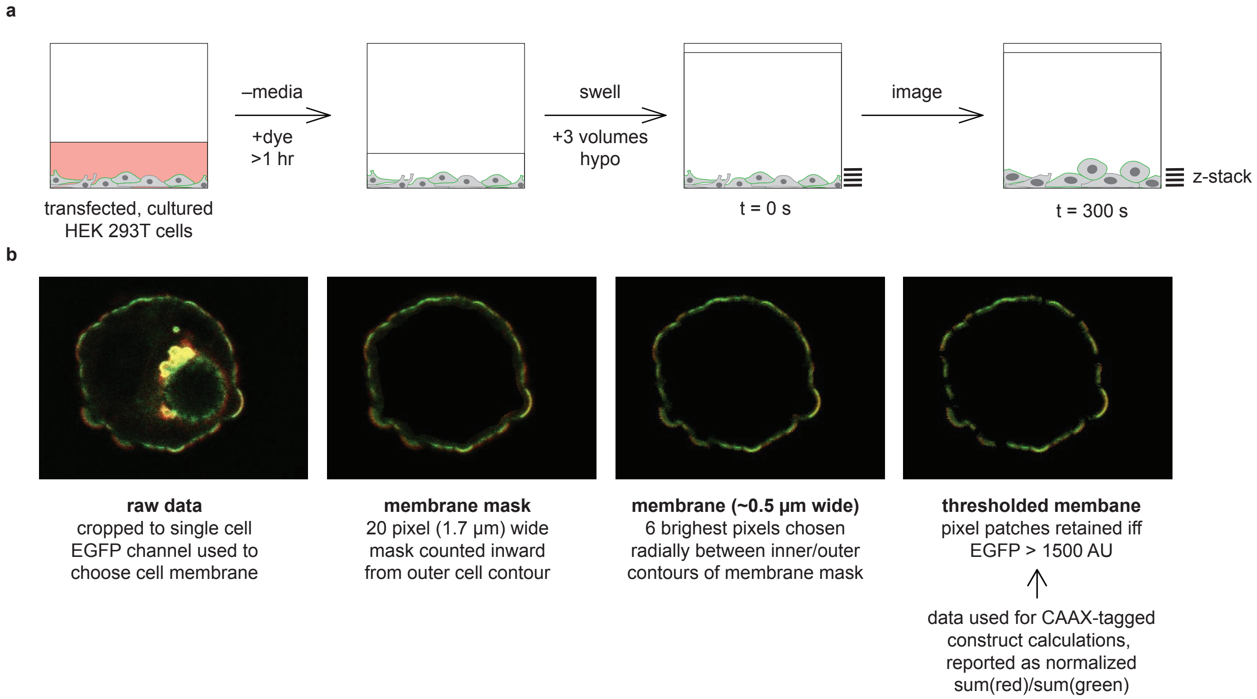


**Supplementary Figure 1. TRAAK(cp/sp)HTx confocal imaging and analysis.** (**a**) Experiment setup of TRAAK (cp/sp)HTx imaging screen. HEK293T cells transfected with construct were treated with 0.5 µM dye for at least one hour prior to imaging. Swelling was induced via the addition of three volumes of hypotonic solution, after which a 4 slice, 10 µm tall z-stack was collected every 30 seconds for 5 minutes. (**b**) Data workup pipeline. Individual fluorescent cells and indistinguishable cell patches were cropped to separate files via deletion/omission of the EGFP signal, then masked according to the EGFP signal in the plasma membrane (see methods). Within the resultant 20-pixel wide membrane mask, the brightest 6 adjacent pixels ("pixel patch") were chosen along each of a series of lines plotted radially from the inner to outer mask contour to produce an approximately 0.5 µm wide cell membrane. The final thresholded membrane data used in all TRAAK(cp/sp)HTx figures/calculations comprises only pixel patches of average EGFP signal greater than 1500 AU.

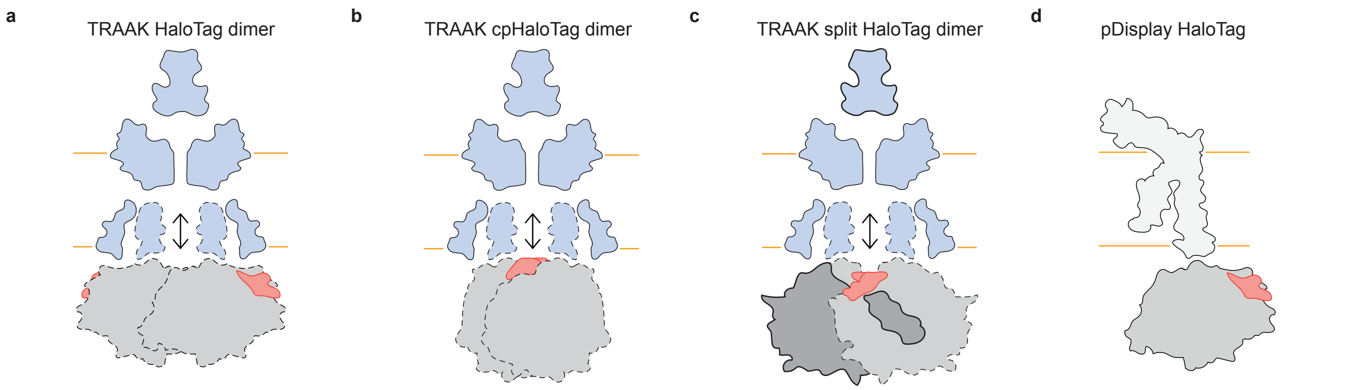


**Supplementary Figure 2. Cartoon depictions of TRAAK (cp/sp)HaloTag based on AlphaFold-predicted structures.**

(**a**–**d**) Cartoon depictions of TRAAK HaloTag, TRAAK cpHaloTag, TRAAK spHaloTag, and pDisplay HaloTag protein, respectively, derived from AlphaFold-predicted structures. HT = HaloTag, cp = circularly permuted, sp = split, Ctrm = TRAAK C-terminus, x = CAAX. Dotted lines denote TM4–HaloTag linkage and associated movement. Bolded lines in **c** denote additional linkage between TM1 (out of plane of page) and N-terminal spHaloTag, which is relatively immobile during TRAAK opening. TRAAK, blue; HaloTag, grey; fluorogenic dye, red.

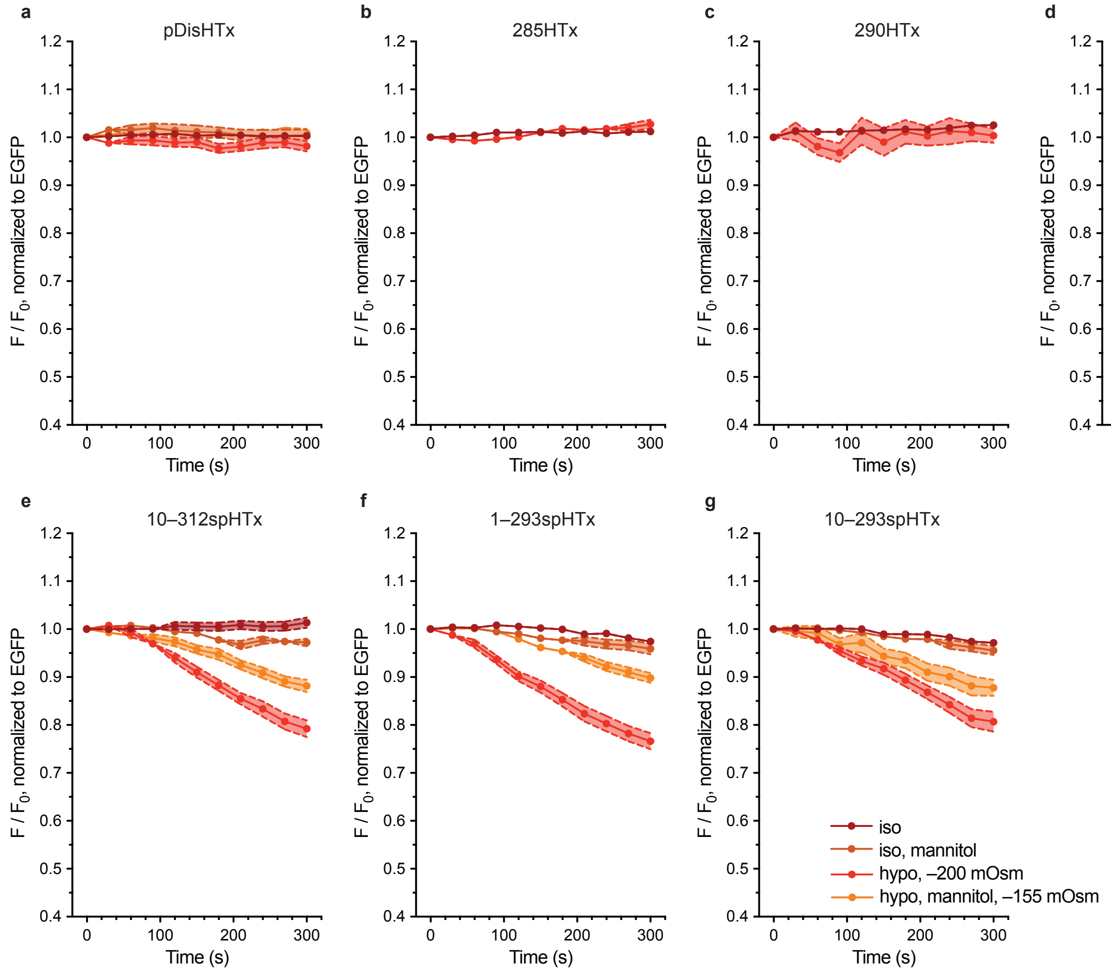


**Supplementary Figure 3. TRAAK(cp/sp)HTx – $JF_{585}$ screen.**

(**a**–**g**) Change in brightness of $JF_{585}$ relative to EGFP over time in membranes of HEK293T cells transiently transfected with pDisHTx, 285HTx, 290HTx, 290cpHTx, 1–312spHTx, 1–293spHTx, or 10–293spHTx, respectively. Iso/hypo: isotonic/hypotonic solution added. Mannitol = mannitol-based solutions. See Methods for solution composition. Data shown as mean ± s.e.m.

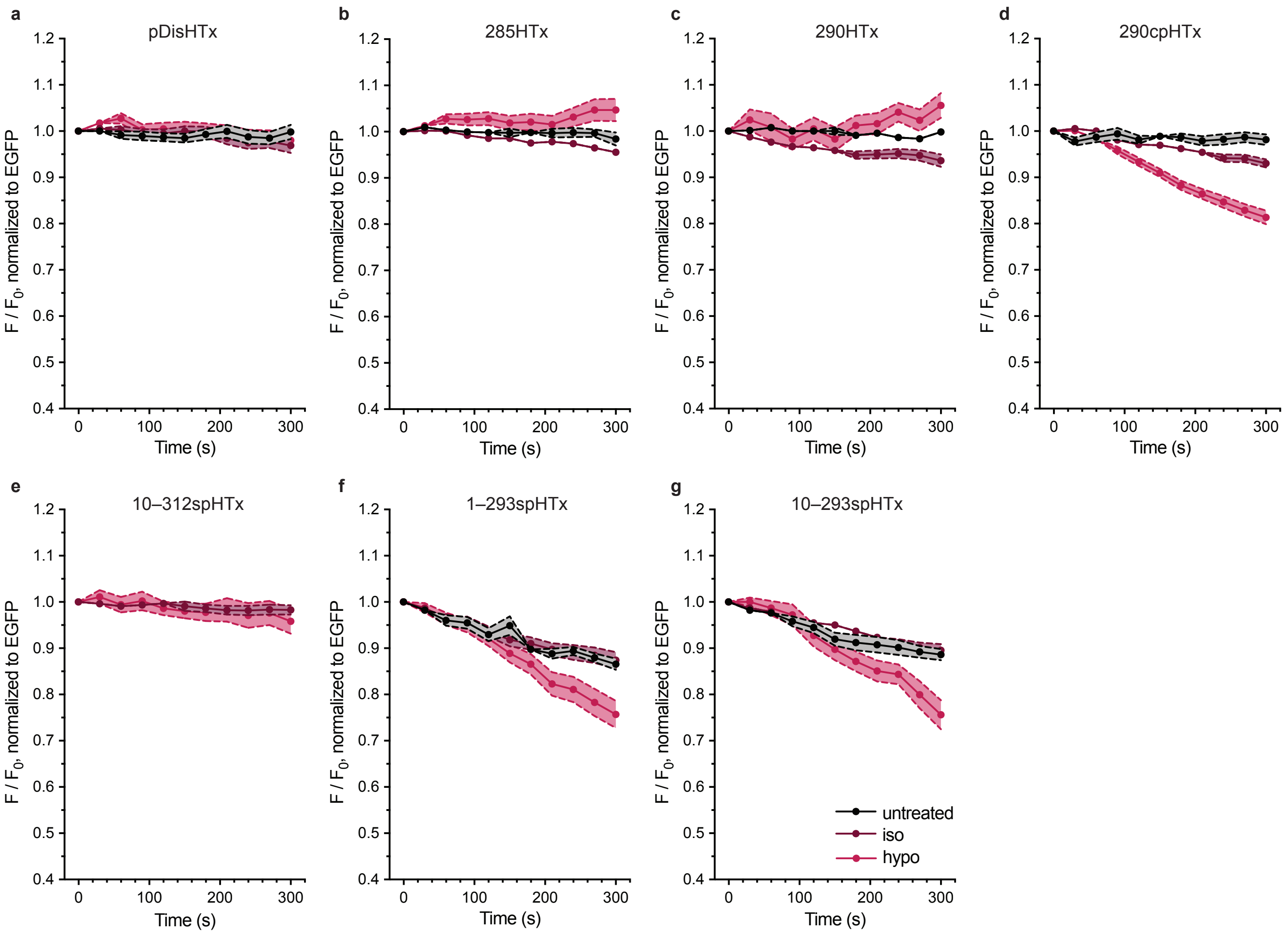


**Supplementary Figure 4. TRAAK(cp/sp)HTx – $JF_{635}$ screen.**
(**a**–**g**) Change in brightness of $JF_{635}$ relative to EGFP over time in membranes of HEK293T cells transiently transfected with pDisHTx, 285HTx, 290HTx, 290cpHTx, 1–312spHTx, 1–293spHTx, or 10–293spHTx, respectively. Untreated: no solution added; iso/hypo: three volumes of isotonic/hypotonic solution added <5 seconds prior to t = 0 s z-stack, respectively. Aligned to match **Supplementary Figures 3**. Data shown as mean ± s.e.m.

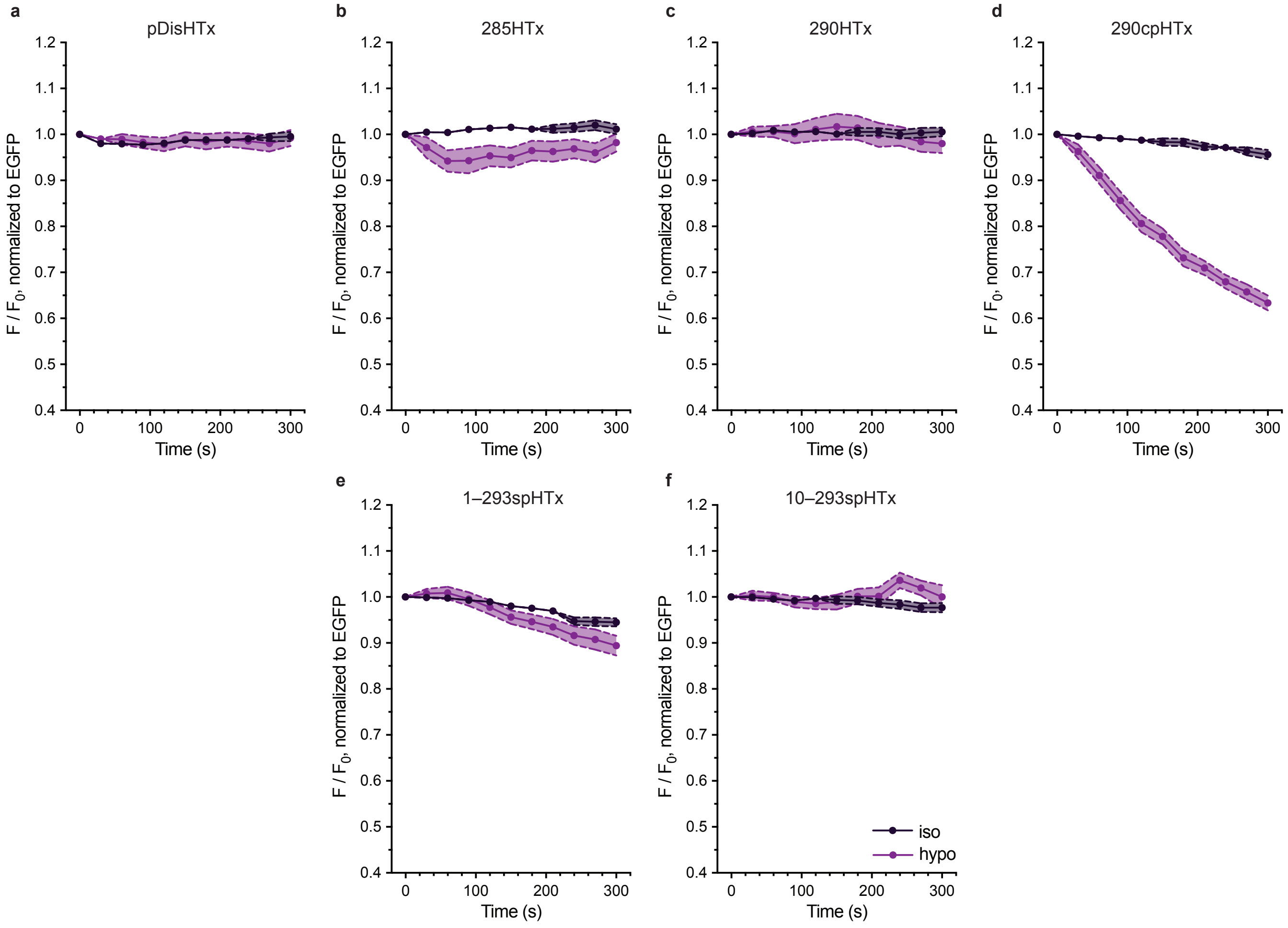


**Supplementary Figure 5. TRAAK(cp/sp)HTx – BF$_{646}$ screen.**

(**a**–**f**) Change in brightness of BF$_{646}$ relative to EGFP over time in membranes of HEK293T cells transiently transfected with pDisHTx, 285HTx, 290HTx, 290cpHTx, 1–293spHTx, or 10–293spHTx, respectively. Iso/hypo: isotonic/hypotonic solution added. Aligned to match **Supplementary Figures 3 and 4**. Data shown as mean ± s.e.m.

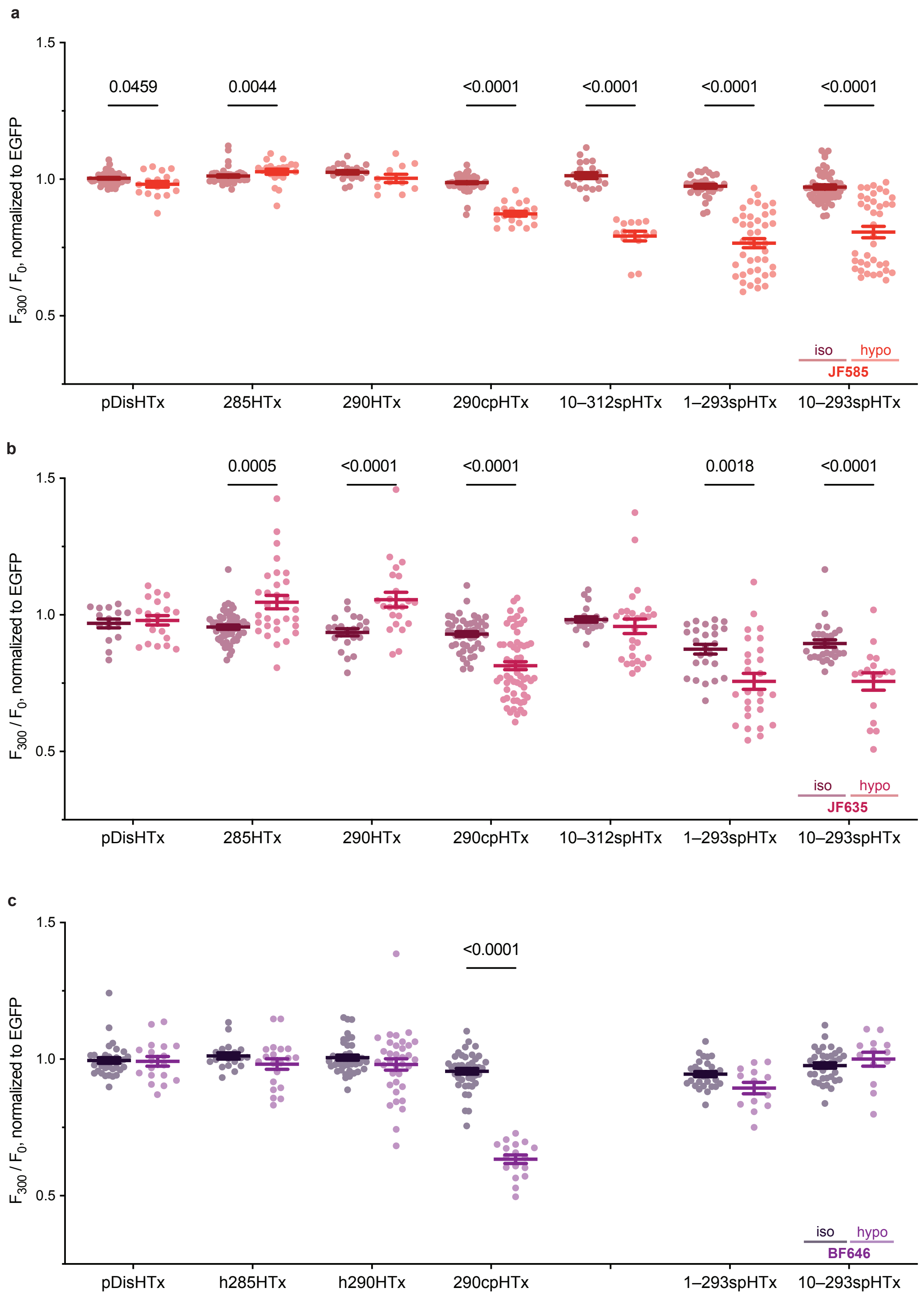


**Supplementary Figure 6. Summary of TRAAK(cp/sp)HTx screens (t = 300 s data from Supplementary Figures 3–5).**

Change in brightness of (**a**) $JF_{585}$, (**b**) $JF_{635}$, or (**c**) $BF_{646}$ relative to EGFP after 5 minutes in membranes of HEK293T cells transiently transfected with described constructs. Iso/hypo: isotonic/hypotonic solution added. Data shown as mean ± s.e.m. p values calculated from multiple unpaired Mann-Whitney tests with no correction for multiple comparisons. p values $< 0.05$ depicted above related data sets.

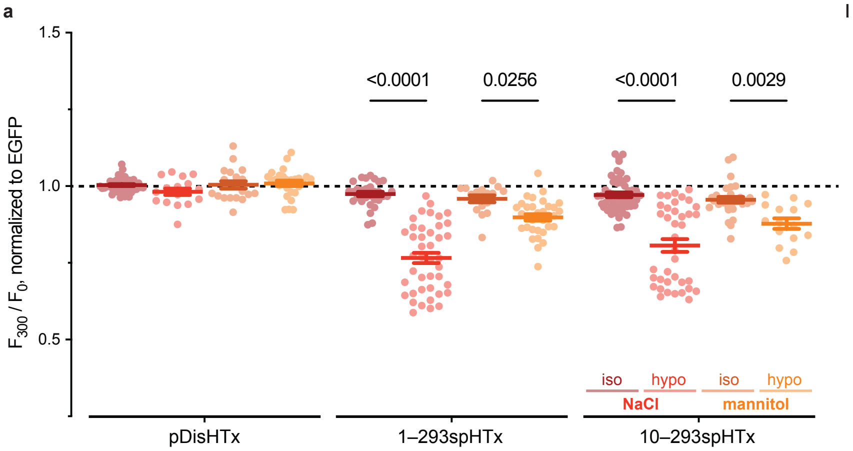


**Supplementary Figure 7: TRAAK spHaloTags respond to osmolarity change, not specific osmolyte.** Change in brightness of $JF_{585}$ relative to EGFP after 5 minutes in membranes of HEK293T cells transiently transfected with described constructs. Iso/hypo: isotonic/hypotonic solution added. NaCl = NaCl-based solutions (iso/hypo elsewhere); Mannitol = mannitol-based solutions. See Methods for solution composition. Data shown as mean ± s.e.m. with individual n's as circles. $n \geq 15$. p values calculated from two-way ANOVA with Tukey's correction; values $< 0.05$ depicted above related data.

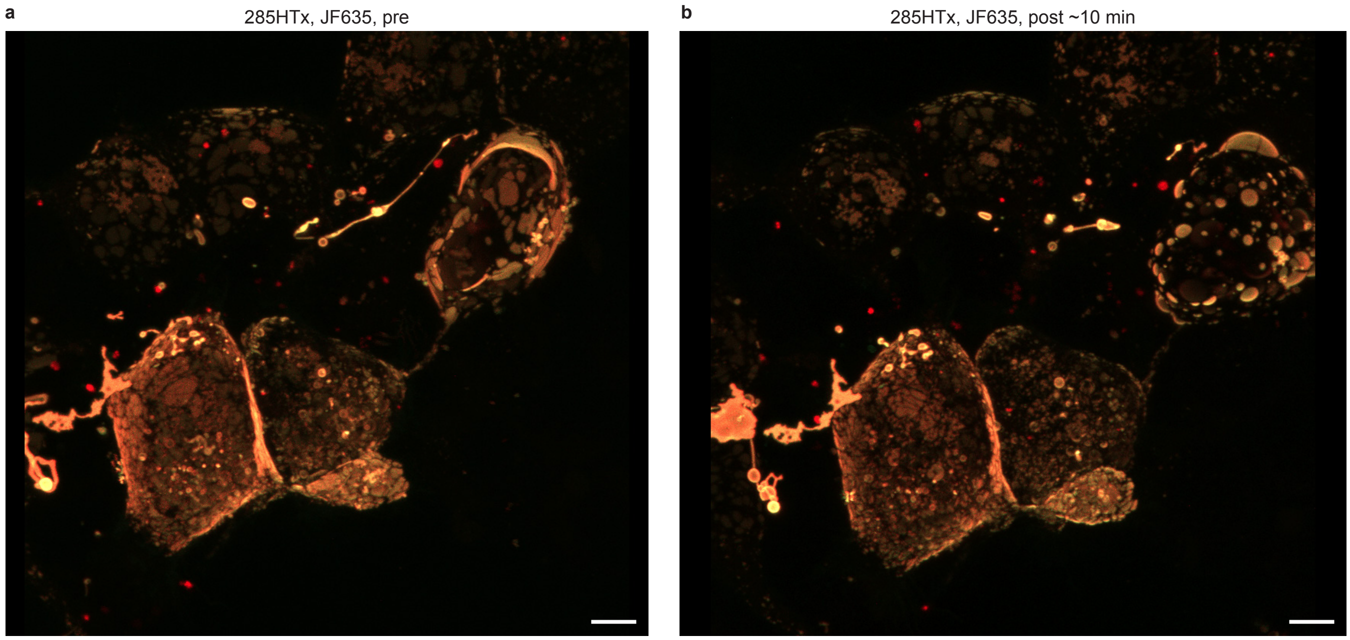


**Supplementary Figure 8. CAAX motif-containing TRAAK(cp/sp)HTx constructs self-sort into raft-like domains in HEK293T plasma membranes.**
z-stacked derived 3D reconstruction of HEK293T cells expressing 285HTx with EGFP signal depicted in green and $JF_{635}$ signal in red (**a**) before (pre) and (**b**) after (post) swelling with the addition of three volumes of hypotonic solution (–200 mOsm). Patches of bright (285HTx-expressing) membrane observed in (**a**) bulge asymmetrically outward upon swell in (**b**) compared to adjacent nonfluorescent membranes. Distance scale bar = 5 µm.

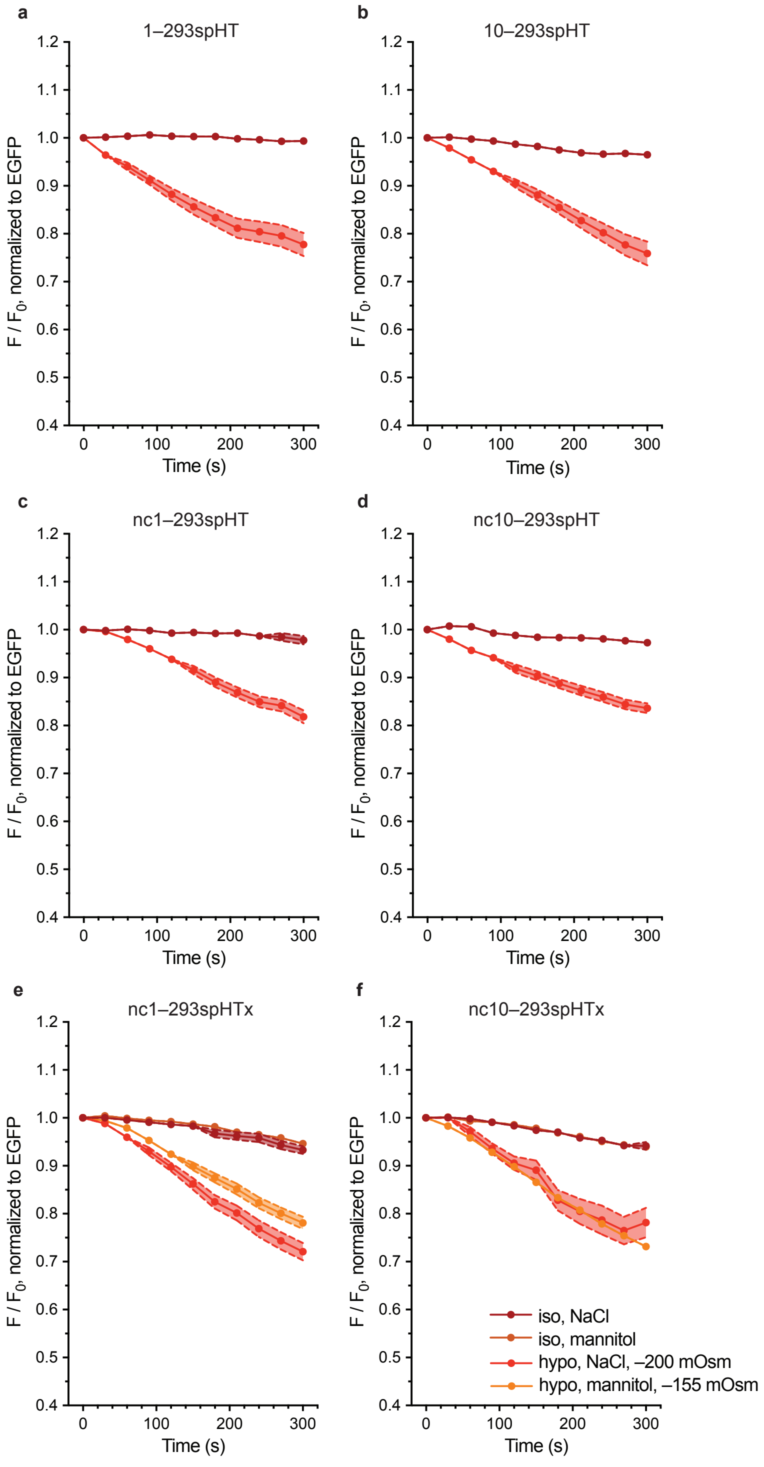


**Supplementary Figure 9. Nonconducting TRAAKspHT(x) – $JF_{585}$ screen.**
(**a**–**f**) Change in brightness of $JF_{585}$ relative to EGFP over time in membranes of HEK293T cells transiently transfected with 1–293spHT, 10–293spHT, nc1–293spHT, nc10–293spHT, nc1–293spHTx, or nc10–293spHTx, respectively. Iso/hypo: isotonic/hypotonic solution added. NaCl = NaCl-based solutions; Mannitol = mannitol-based solutions. See Methods for solution composition. Data shown as mean ± s.e.m. See **Supplementary Figure 3f–g** for comparison.

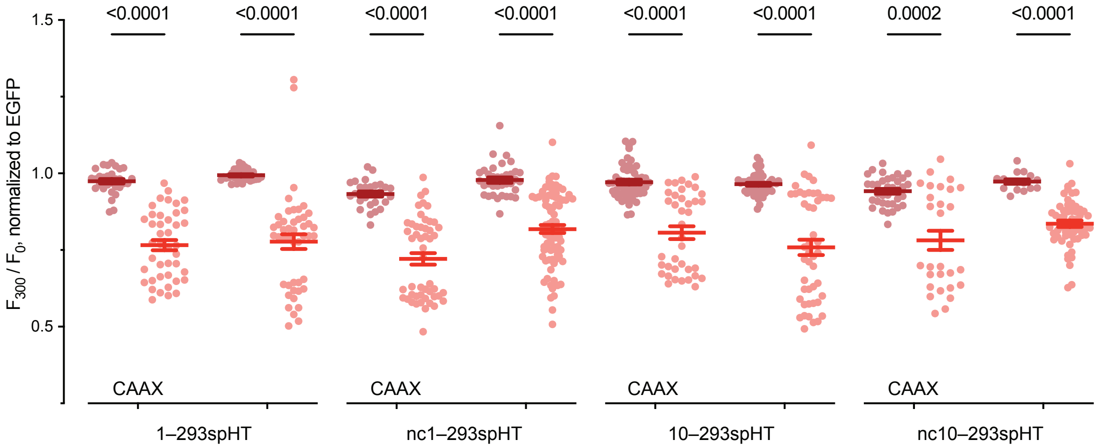


**Supplementary Figure 10. Neither CAAX tag nor potassium conduction are required for sensor function.**

Change in brightness of $JF_{585}$ relative to EGFP after 5 minutes in membranes of HEK293T cells transiently transfected with described constructs. Iso/hypo: isotonic/hypotonic solution added. Data shown as mean ± s.e.m. p values calculated from multiple unpaired Mann-Whitney tests with no correction for multiple comparisons.

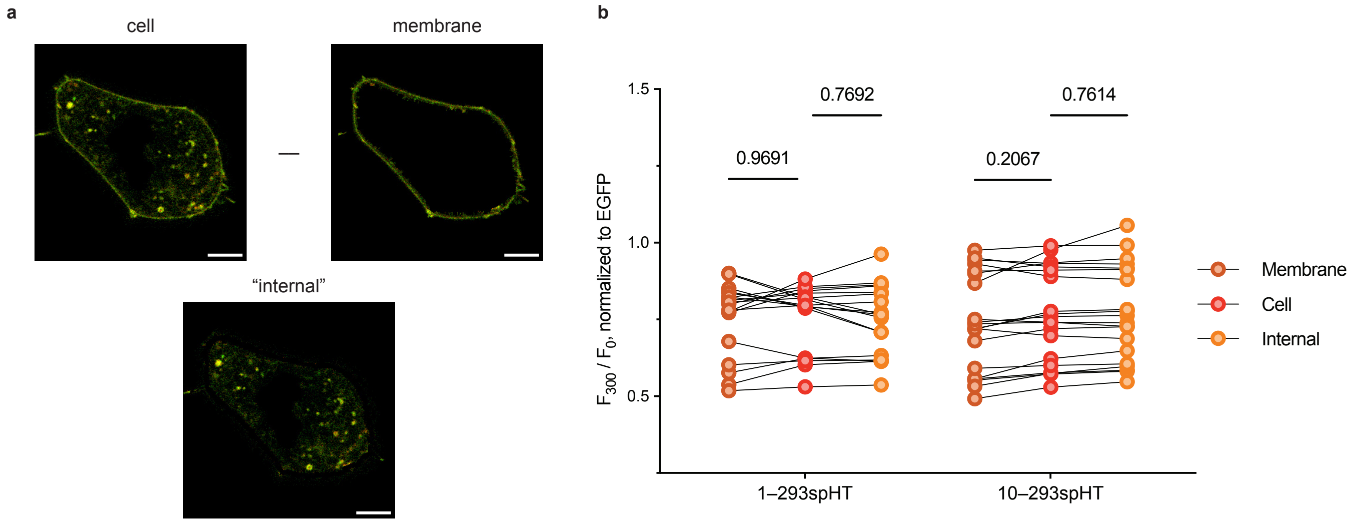


**Supplementary Figure 11. TRAAKspHTs dim to similar degrees in internal and plasma membranes 5 minutes after hypotonic shock.**

(**a**) Data workup pipeline for TRAAKspHT constructs. Membrane data calculated as described in **Supplementary Figure 2b** (thresholded membrane). Cell data derived from cell mask subjected to 1500 AU threshold in the EGFP channel. "Internal" membranes approximated as Cell – Membrane. Distance scale bar = 5 µm. (**b**) Change in brightness of $JF_{585}$ relative to EGFP in HEK293T cells transiently transfected with described constructs after 5 minutes of hypotonic shock (–200 mOsm). Matched membrane, cell, and internal data appear from left to right, calculated as described in **a**. $n \geq 18$ shown as individual circles. p values calculated with two-way ANOVA with Tukey's correction, depicted above related data.

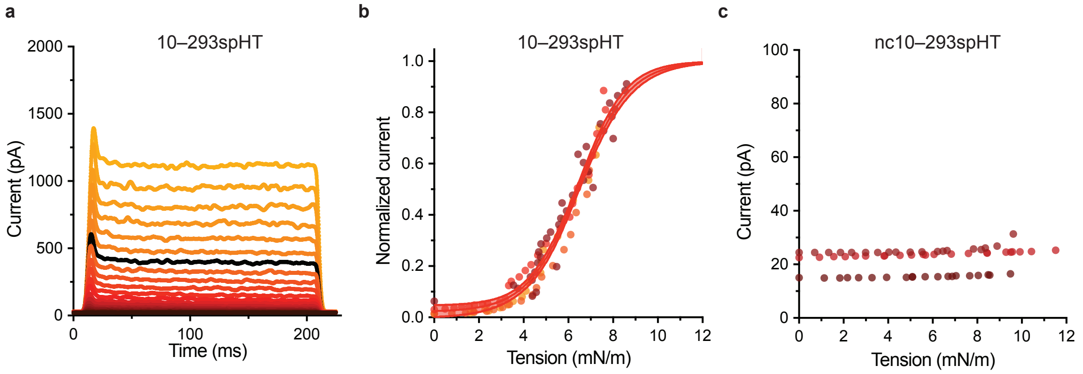


**Supplementary Figure 12. 10–293spHT has a right-shifted tension activation curves.** (**a**) Representative currents produced by 10–293spHT in excised inside-out HEK293T cell patch in response to increasingly negative (in increments of 5 mmHg) pressure steps. Currents elicited by a –70 mmHg pressure step highlighted in black. (**b**) Tension activation curve calculated for 10–293spHT (n = 4) according to best Boltzmann sigmoidal fit (mean ± s.e.m., solid lines). Normalized data from individual inside-out patches shown as circles, colored by cell. (**c**) Tension conduction plot for nc10–293spHT (n = 3). Normalized data from individual inside-out patches shown as circles, colored by cell. nc1–293 data is flat and could neither be normalized nor fit to a Boltzmann sigmoidal curve.

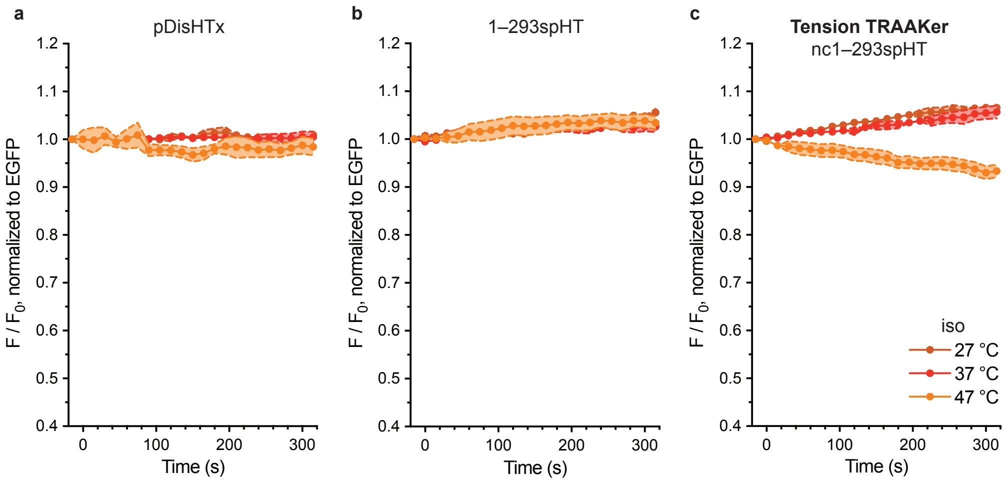


**Supplementary Figure 13. (nc)1–293spHT brightness remains relatively stable under isotonic conditions.**
(**a**–**c**) Change in brightness of $JF_{585}$ relative to EGFP over time in membranes of HEK293T cells transiently transfected with pDisHTx, 1–293spHT, or Tension TRAAKer, respectively, at the described temperatures. Iso: three volumes of isotonic solution added <2 seconds prior to t = 0 s image. Data shown as mean ± s.e.m. $n \geq 7$.

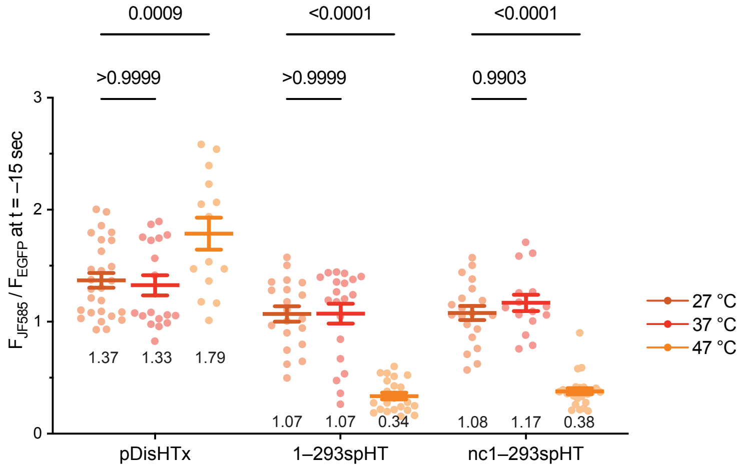


**Supplementary Figure 14. (nc)1–293spHT is temperature sensitive.**
$JF_{585}$/EGFP at t = –15 seconds (post equilibration, pre-experiment start) in pDisHTx-, 1–293spHT-, and nc1–293spHT-expressing HEK293T cells at listed temperatures. p values calculated from two-way ANOVA with Tukey's correction; depicted above related data. Mean $F_{JF585}/F_{EGFP}$ values depicted below related data.

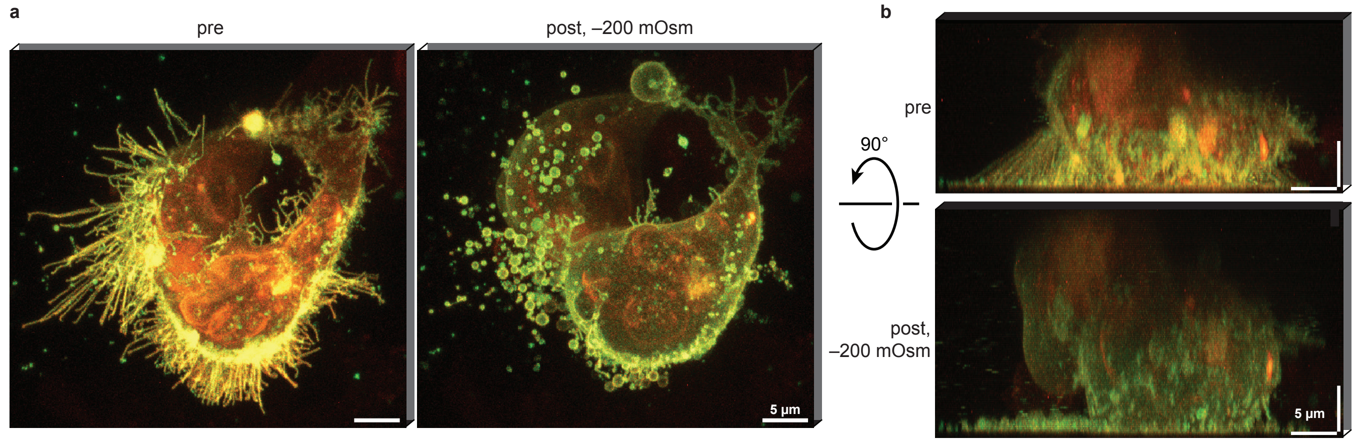


**Supplementary Figure 15. Tension TRAAKer operates in three dimensions.**
(**a**–**b**) Representative three-dimensional reconstruction of Tension TRAAKer-expressing cell pre- and post-hypotonic shock depicted from orthogonal viewpoints. Brightness scaling maintained within same channel and cell across images. Distance scale bar = 5 µm.

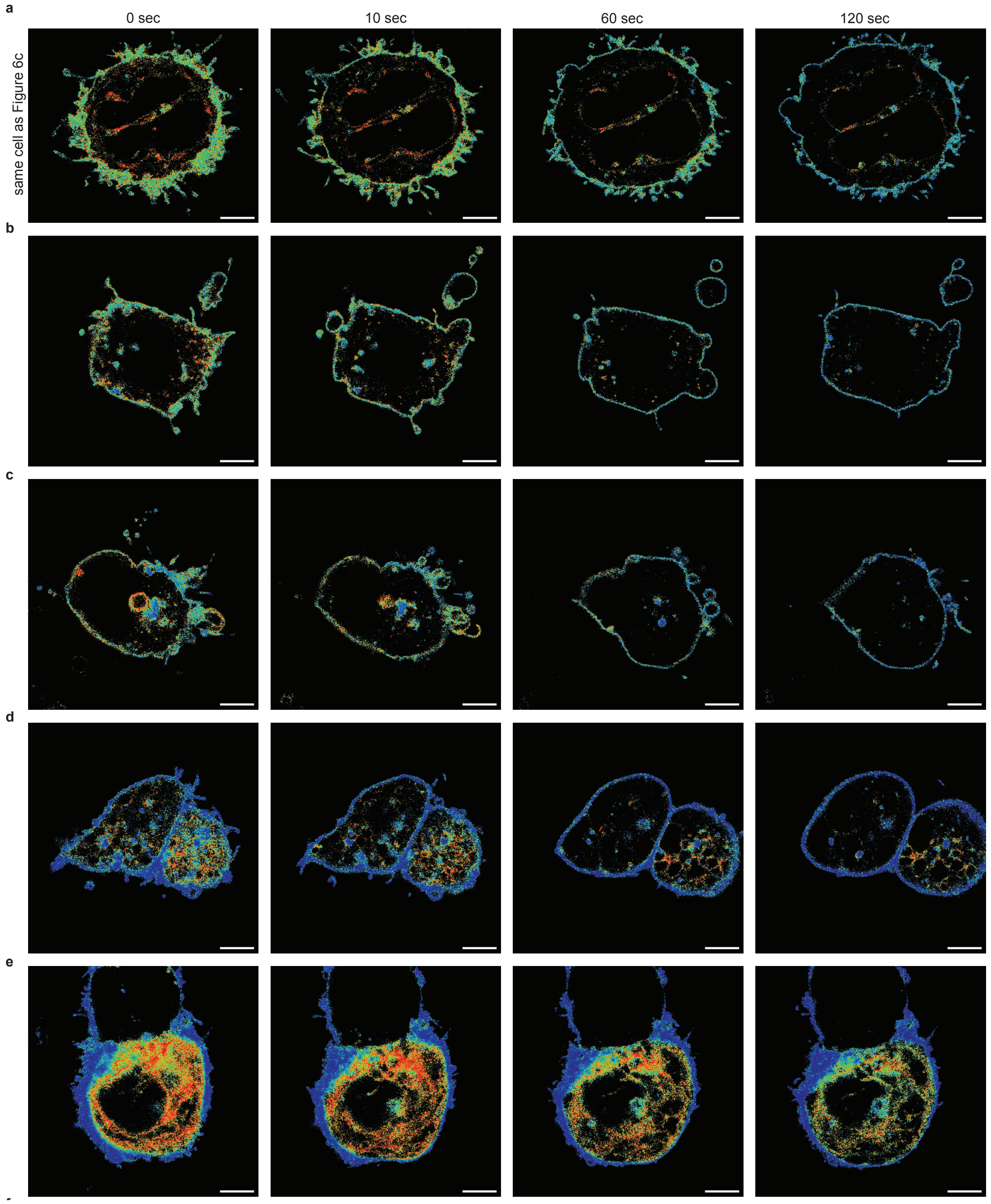


144

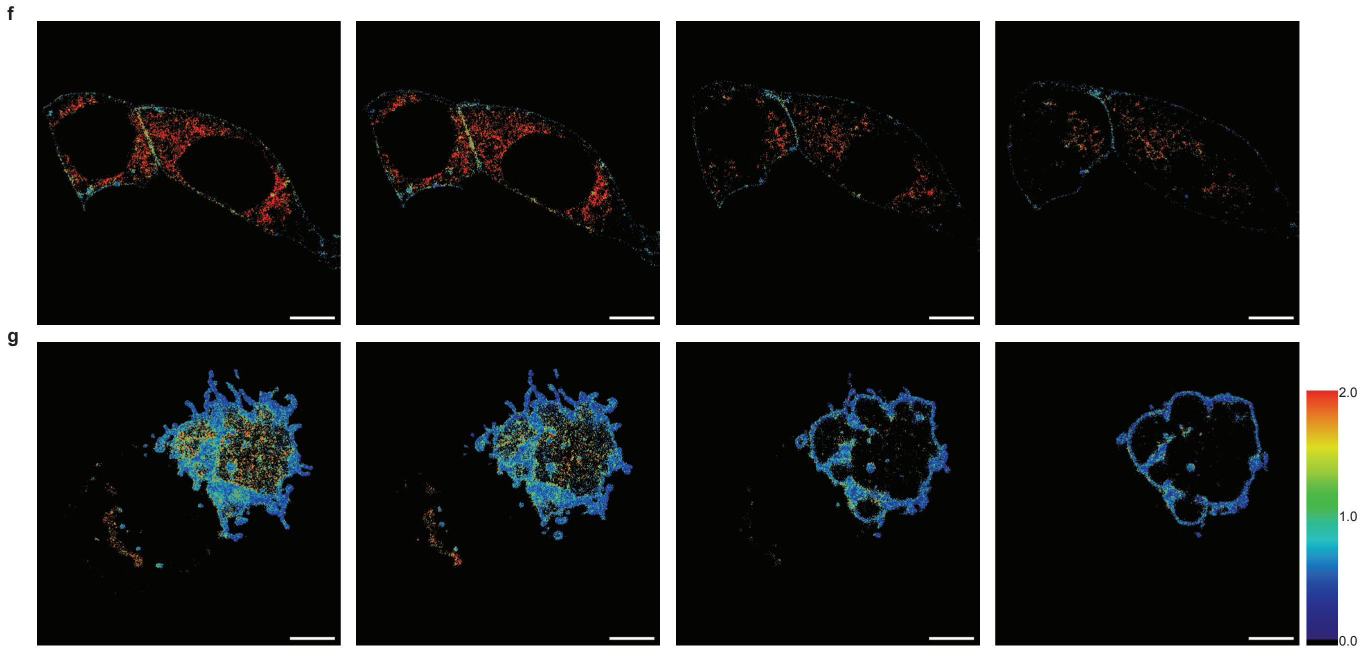


**Supplementary Figure 16. Tension TRAAKer responds to hypotonic shock with subcellular variability.**
(**a–g**) Tension TRAAKer cells at 37 °C at listed time points post-hypotonic (–200 mOsm) shock. Color coded by $JF_{585}$/EGFP pixel value.

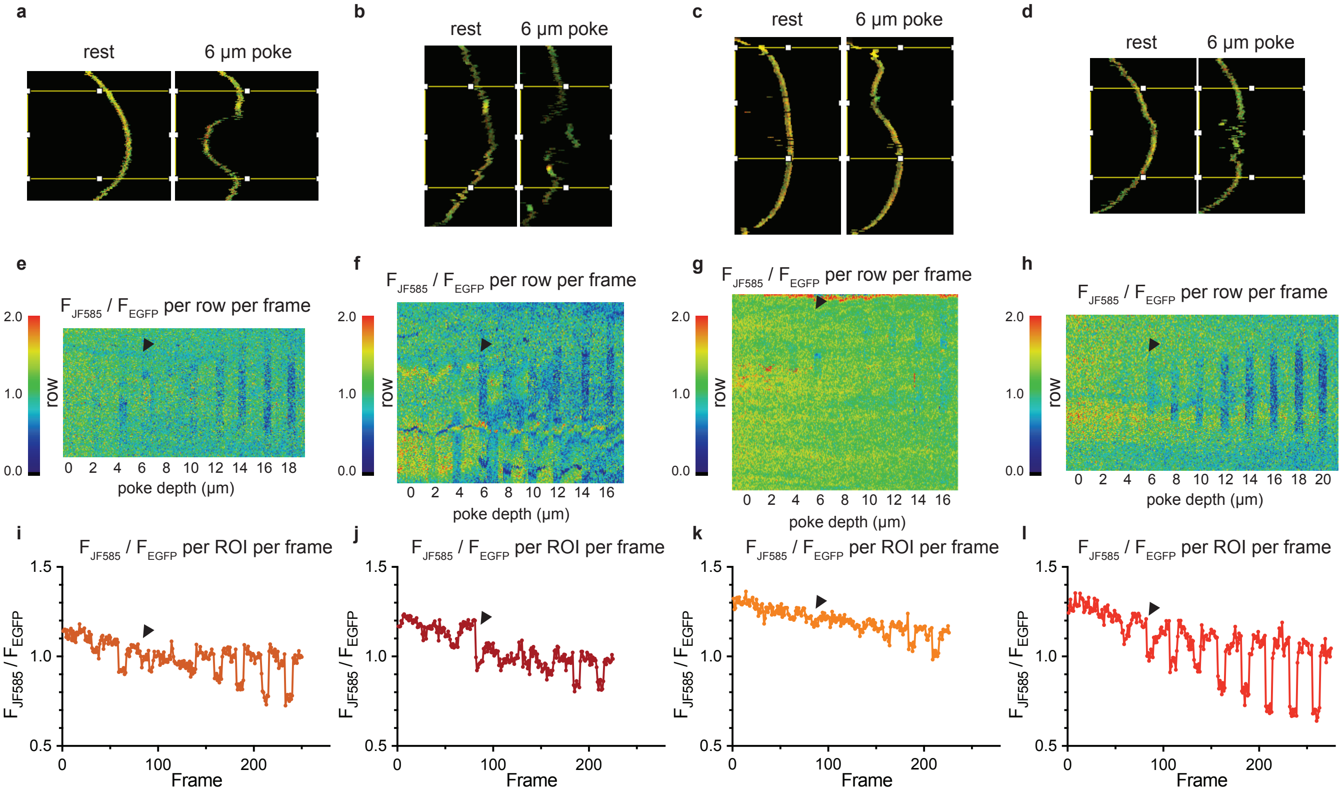


**Supplementary Figure 17. Tension TRAAKer dims in response to cell poking.**

(**a**–**d**) Representative images of masked membranes calculated from cell poking assay for four different cells (**d** = representative cell depicted in **Figure 6**). $JF_{585}$ depicted in red, EGFP in green. ROI used for data workup in **i**–**l** highlighted in yellow box. (**e**–**h**) Kymographs of Tension TRAAKer cells subjected to 5-second-long cell pokes of increasing depth from left to right. Each column is a laterally compressed membrane mask of a single frame of video, constructed from the average $JF_{585}$/EGFP value per masked row of pixels. Color coded by $JF_{585}$/EGFP value, matched to **a**–**d**. (**i**–**l**) Plots of Tension TRAAKer response to pokes of increasing depth within ROIs depicted in **a**–**d** (encompassing, approximately, the length of membrane affected by the deepest cell poke), calculated as total $JF_{585}$ fluorescence divided by total EGFP fluorescence. x-axis (frame) scaled to kymograph above each plot. Color coded according to **Figure 6h**. Black arrows identify location of 6 µm poke in all plots for reference.

#### 2. Supplementary Tables

**Supplementary Table 1.** Summary of published fluorogenic dye properties.[1,2,3]

| Dye | $\lambda_{ex}$ (nm) | $\lambda_{em}$ (nm) | ε ($M^{-1}cm^{-1}$) | φ | εφ ($M^{-1}cm^{-1}$) | ΔF/F* |
|---|---|---|---|---|---|---|
| **BF646** | 646 | 662 | 66,000 | 0.29 | 19,140 | 23.1 |
| **JF635** | 635 | 652 | 167,000 | 0.56 | 93,520 | 66 |
| **JF585** | 585 | 609 | 156,000 | 0.78 | 121,680 | 98 |
| **EGFP** | 488 | 507 | 55,900 | 0.60 | 33,540 | — |

*Reported for HaloTag ligand; remaining parameters reported for free dye.
**Calculated from $F/F_0$ = 24.1.

**Supplementary Table 2.** Summary of **Figure 3** TRAAK(spHT) tension activation curves.

| Construct | $T_{50}$ (mN/m) | Slope (m/mN) | $R^2$ | n |
|---|---|---|---|---|
| **TRAAK–EGFP** | 3.50 ± 0.16 | 1.08 ± 0.12 | 0.8956 | 5 |
| **1–293spHT** | 6.65 ± 0.07 | 1.53 ± 0.07 | 0.9345 | 8 |
| **10–293spHT** | 6.47 ± 0.05 | 1.13 ± 0.05 | 0.9620 | 4 |

All mean ± s.e.m.

**Supplementary Table 3.** Summary of **Figure 4** 47 °C (nc)1–293spHT exponential fits.

| Construct | Temp. | Condition | $Y_0$ | Plateau | Half Life (s) | $R^2$ | n |
|---|---|---|---|---|---|---|---|
| 1–293spHT | 47°C | hypo, 5 min | 0.93 ± 0.02 | 0.51 ± 0.01 | 48.6 | 0.5351 | 17 |
| nc1–293spHT | 47°C | hypo, 5 min | 0.92 ± 0.01 | 0.45 ± 0.01 | 46.9 | 0.6747 | 16 |

All mean ± s.e.m.

**Supplementary Table 4.** Summary of **Figure 5** (nc)1–293spHT hypotonic shock exponential fits.

| Construct | Temp. | Condition | $Y_0$ | Plateau | τ (s) | $R^2$ | n |
|---|---|---|---|---|---|---|---|
| 1–293spHT | 37 °C | hypo, 10 min | 0.98 ± 0.01 | 0.56 ± 0.01 | 162.9 | 0.7915 | 10 |
| 1–293spHT | 37 °C | hypo, 5 min rec | 0.98 ± 0.01 | 0.54 ± 0.06 | 209.3 | 0.6537 | 10 |
| 1–293spHT | 37 °C | hypo, 2 min rec | 0.97 ± 0.01 | 0.66 ± 0.11 | 138.7 | 0.7378 | 8 |
| nc1–293spHT | 37 °C | hypo, 10 min | 0.95 ± 0.01 | 0.47 ± 0.02 | 263.2 | 0.7229 | 12 |
| nc1–293spHT | 37 °C | hypo, 5 min rec | 0.97 ± 0.02 | 0.56 ± 0.09 | 203.4 | 0.3867 | 11 |
| nc1–293spHT | 37 °C | hypo, 2 min rec | 0.95 ± 0.02 | 0.58 ± 0.58 | 204.0 | 0.2849 | 8 |

All mean ± s.e.m.

**Supplementary Table 5.** Summary of **Figure 5** (nc)1–293spHT recovery exponential fits.

| Construct | Temp. | Condition | $Y_0$ | Plateau | τ (s) | $R^2$ | n |
|---|---|---|---|---|---|---|---|
| 1–293spHT | 37 °C | hypo, 5 min rec | 0.10 ± 0.80 | 0.88 ± 0.13 | 264.21 | 0.1885 | 10 |
| 1–293spHT | 37 °C | hypo, 2 min rec | 0.33 ± 0.15 | 1.24 ± 0.06 | 266.7 | 0.5184 | 8 |
| nc1–293spHT | 37 °C | hypo, 5 min rec | –0.44 ± 1.66 | 1.01 ± 0.10 | 197.9 | 0.1895 | 11 |
| nc1–293spHT | 37 °C | hypo, 2 min rec | 0.52 ± 0.11 | 1.21 ± 0.04 | 252.5 | 0.5441 | 8 |

All mean ± s.e.m.
